# Aneuploidy, polyploidy, and loss of heterozygosity distinguish serial bloodstream isolates of *Candida albicans*

**DOI:** 10.1101/2025.10.27.684856

**Authors:** Nancy E. Scott, Xin Zhou, Elizabeth Wash, Christopher Zajac, Susan E. Kline, Serin E Erayil, Anna Selmecki

**Affiliations:** University of Minnesota, Bioinformatics and Computational Biology Program; University of Minnesota, Department of Microbiology and Immunology; University of Minnesota, Molecular, Cellular, Developmental Biology and Genetics Program; University of Minnesota, Department of Medicine, Division of Infectious Diseases and International Medicine

**Keywords:** *Candida albicans*, antifungal drug tolerance, *ERG251*, polyploidy, serial isolates, within-host diversity

## Abstract

**Background:** The opportunistic pathogen *Candida albicans* is the leading species causing invasive *Candida* infections worldwide. Genomic variation is widespread in clinical isolates and complicates identification of genetic variants underlying antifungal drug resistance and tolerance. Our understanding of genomic and phenotypic diversity during invasive infections is limited and studies of serial isolates from individual patients are uncommon. We performed comparative analyses of 101 *C. albicans* bloodstream isolates from 55 patients in the Minnesota Minneapolis-Saint Paul region, including serial isolates from 19 patients. We analyzed the phylogenetic relationships of these isolates relative to 199 globally-collected public *C. albicans* genomes.

**Results:** This study’s regional isolates span the phylogenetic diversity of *C. albicans*; 6 isolates represent novel outliers to known clades. Serial isolates from individual patients were separated by limited single nucleotide polymorphisms. Nevertheless, we identified extensive large-scale genomic variation between serial isolates including polyploidy, aneuploidy, copy number variation, loss of heterozygosity, and chromosomal rearrangements. We demonstrated how a heterozygous *ERG251* loss of function variant drives azole tolerance in a clinical isolate from a patient with a history of recurrent infections. Using serial isolates, we demonstrated that polyploidy provides an adaptive advantage in the presence of fluconazole despite the absence of overt antifungal drug resistance.

**Conclusions:** Our analysis of serial isolates reveals the genomic plasticity of *C. albicans* during invasive infections and identifies variation driving antifungal drug tolerance. Our findings reveal limitations in current antifungal susceptibility testing and highlight the need to account for genomic and phenotypic variation during invasive *Candida* infections.

## Background

Invasive *Candida* infections represent a significant and growing threat to global public health, particularly among immunocompromised individuals (1–3). Invasive *Candida* infections, including candidemia, are associated with high morbidity and mortality between 27% and 72% (2,4,5). *Candida albicans* is the leading cause of invasive infections worldwide and its ability to transition from commensal organism to infectious pathogen underlies its importance in settings involving broad-spectrum antibiotics, invasive medical devices such as central venous catheters, and immunosuppression (1–3,5). While genomic studies have advanced our understanding of *C. albicans* biology, limitations remain in translating these findings into clinical practice, including linking genotypes to phenotypes and integrating these results into real-time treatment strategies.

At least twenty-three major phylogenetic clusters (also referred to as clades) of *C. albicans* have been identified through multilocus sequence typing (MLST) and whole genome sequencing (WGS)-based analyses of clinical, commensal and environmental strains (6–9). These phylogenetic analyses have also identified isolates that do not group with known clusters, suggesting that existing clusters may need to be further refined or that rare clusters still have not been well characterized (7,8). How much regional geographic variation exists in the prevalence of *C. albicans* phylogenetic clusters, and what the clinical significance may be, is not known.

*C. albicans* is a heterozygous diploid yeast and well known for its genomic plasticity: large scale genomic alterations such as gross chromosomal rearrangements, aneuploidy (whole chromosome copy number change), loss of heterozygosity (LOH) and copy number variation (CNV, defined herein as impacting the copy number of a segment of chromosome ≥ 10 kb) (10–17). The frequency and effect of many of these genomic changes on adaptation and fitness have been studied both *in vitro* and *in vivo* using experimental evolution (16–22). Aneuploidy, CNVs, LOH and karyotypic changes are also frequently found in clinical isolates (13,23–25). However, only limited data are available about the spectrum and frequency of genomic alterations within an individual patient or during ongoing infection.

Comparative genomic analysis of serial clinical isolates of *Candida* is uncommon and has typically focused on chronic infections, with samples collected months to years apart (23,26–28). Current clinical laboratory practices are based on the assumption that invasive infections are clonal populations (29,30). For example, antimicrobial susceptibility testing is performed only on a single colony isolated from a clinical culture, however this technique does not reflect the diversity of an infecting population (31,32). How within-host genomic diversity influences the outcome of invasive infections is poorly understood.

Antifungal drug treatment options for invasive candidiasis are limited to only three major classes: azoles, echinocandins and polyenes (33). Although acquired drug resistance in *C. albicans* bloodstream isolates in the United States is low (34), some susceptible isolates exhibit drug tolerance, defined as persistent growth in drug concentrations above their minimum inhibitory concentration (MIC) (35). Tolerance has been associated with persistent infections and treatment failure in otherwise drug-susceptible isolates (36). Until now, only a few mechanisms driving drug tolerance have been described (37).

We surveyed the species distribution and antifungal susceptibility of 289 *Candida* bloodstream isolates collected in an academic health center and 5 affiliated hospitals in the Minneapolis – Saint Paul region of Minnesota (38). Here, we performed whole genome sequencing (WGS) and comprehensive comparative analysis of the 101 *C. albicans* bloodstream isolates obtained by our survey study, comparing their genotypes and phenotypes within and across patients. We analyzed the phylogenetic structure of our isolate set in the context of 199 publicly available *C. albicans* genomes and identified a heterozygous *ERG251* nonsense variant driving azole tolerance in one clinical isolate. By comparing the karyotypes, ploidy, CNVs, LOHs, and single nucleotide polymorphisms (SNPs) of 19 sets of serial isolates collected from individual patients, we demonstrate extensive within-host genetic variation of infecting populations which may contribute to rapid adaptation and persistent infection even in the absence of drug resistance.

## Results

### Whole genome sequencing and phenotyping of *Candida albicans* bloodstream isolates

101 *C. albicans* isolates were collected from 55 patients as part of a survey of *Candida* bloodstream isolates (38). A “case” refers to all isolates collected from an individual patient. Cases were each assigned a numeric code (case 1, case 2, etc.), and multiple isolates from the same patient were labeled by order of collection (case 1: isolate 1-1, isolate 1-2, etc.). Eighteen patient cases included multiple isolates over time periods of less than 30 days, defined as serial isolate cases. One serial isolate case, and one additional patient case, had multiple isolates collected over more than 100 days apart and these are defined as recurrent cases.

Phenotypic analysis of all isolates was performed as described previously (38). Antifungal drug susceptibility was measured by the EUCAST method of minimum inhibitory concentration (MIC) to fluconazole, micafungin and amphotericin B. All 101 isolates were sensitive to all drugs, however, a wide range of fluconazole tolerance (measured by supra-MIC growth, SMG, see Methods) was observed (Supplemental Table S1). We also determined the 24-hour growth rate and doubling time in rich media as a proxy for isolate fitness in the absence of drug (Supplemental Table S1). The growth rate/hour (r) in rich media ranged from 0.403 to 0.86, with a median value of 0.685.

We performed WGS and variant calling for the 101 isolates relative to the SC5314 A21 reference genome (39). There were a total of 294,938 high-confidence SNPs and 45,338 small insertions and deletions (indels). Relative to the SC5314 reference strain, the number of SNPs per isolate ranged from 10,130 to 98,302 (0.71– 6.88 SNP/kb) and the number of indels ranged from 3,499 to 15,816 (0.24 – 1.11 indels/kb) (Supplemental Table S1). To evaluate strain diversity of our isolates relative to known MLST profiles of *C. albicans* (7,40), we determined the sequence types (STs) of all study isolates (Supplemental Table S1). We identified a total of 47 different STs, including 27 new STs.

### Six isolates from three patient cases do not cluster with any known phylogenetic clades

To determine the phylogenetic relationship of the isolates, we built a maximum-likelihood tree from the 101 isolates from this study and 199 additional publicly available *C. albicans* genomes (Supplemental Table S2) using 265,581 high-confidence SNPs (8,24). We identified a total of 21 clusters, including 18 that have been previously described (Figure 1) (7,8). We identified three distinct clusters that split the previously described MLST cluster 8 and labeled these as 8A, 8B and 8C. Similarly, cluster 4 was separated into 4A and 4B. Serial isolates from individual patients in this study clustered closely together in all cases. Ninety-five of the 101 isolates from this study clustered within known or subdivided clusters A, 1, 2, 3, 4A, 4B, 7, 8B and 8C. Importantly, six isolates from three patient cases could not be assigned to any previously described clusters (cases 41, 60 and 98).

**Figure 1.**
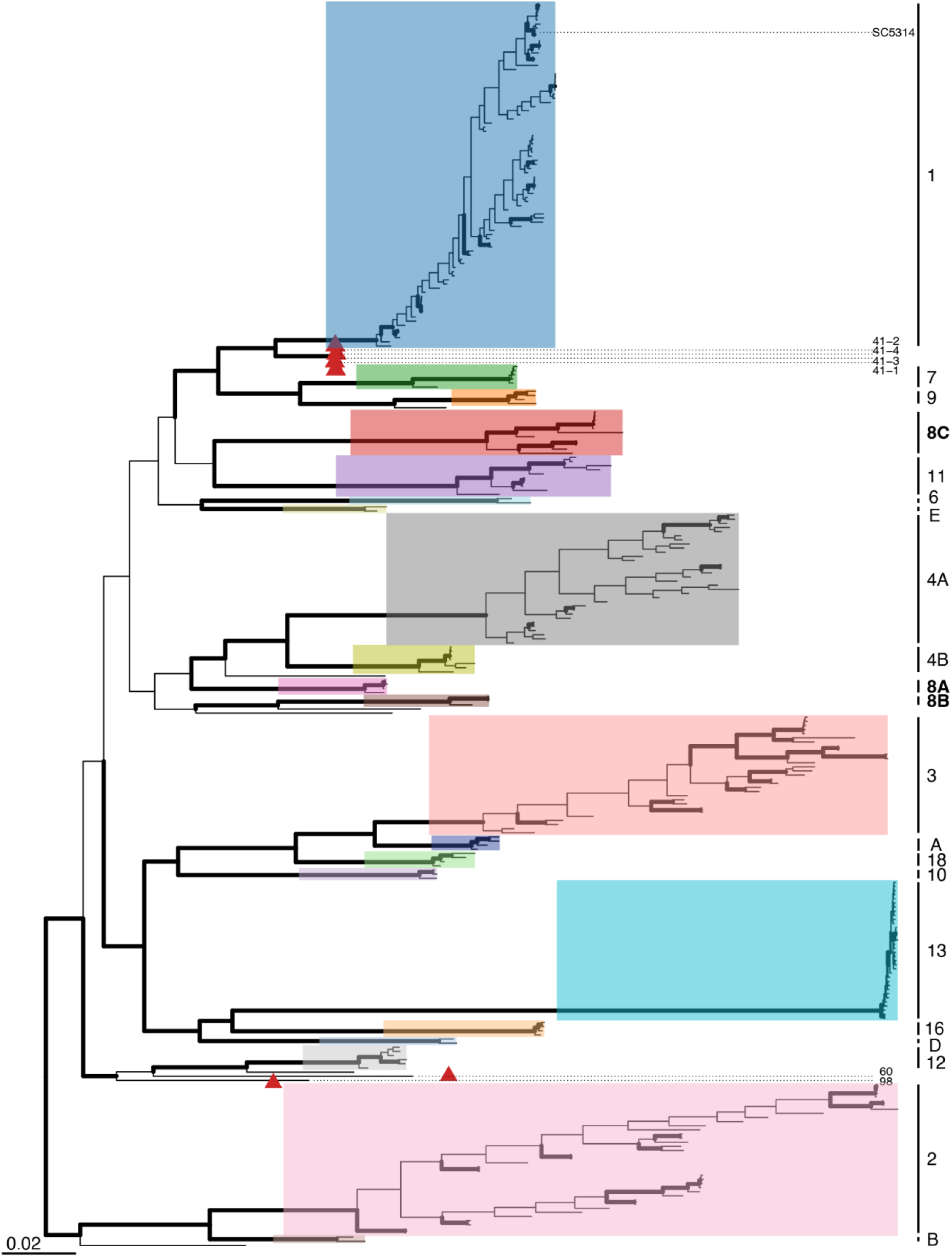
Phylogenetic analysis of 300 isolates identifies 20 major Candida albicans clusters. Maximum-likelihood phylogenetic tree built with 101 new isolates from this study and 199 publicly available genomes (8,24). Cluster labels correspond to Odds et al. (2007) and Ropars et al. (2018). All serial isolates from individual patients in this study cluster together. Six isolates (red triangles) from three cases do not cluster with any previously described clade (patient cases 41, 60 and 98 labeled with dashed lines). MLST clade 4 is divided into 2 clusters designated 4A and 4B by this analysis. MLST clade 8 is divided into 3 clusters designated 8A, 8B and 8C. Bootstrap support > 95% represented by thick, bolded bars. The scale bar represents the number of substitutions per site. Reference isolate SC5314 is indicated. The names of every isolate are detailed in Supplemental Figure S1.

### Loss of heterozygosity patterns are often shared within a phylogenetic cluster

LOH is a major driver of phenotypic diversity in *C. albicans*. To visualize LOH patterns, we plotted the heterozygous SNP density per 5 kb window across the genome for each isolate and ordered isolates by their placement in the phylogenetic tree (Figure 2A). The LOH pattern of individual strains is typically quite unique, akin to a fingerprint, and serial isolates from individual patients have strikingly similar LOH patterns (Figure 2A and Supplemental Figure S2).

**Figure 2.**
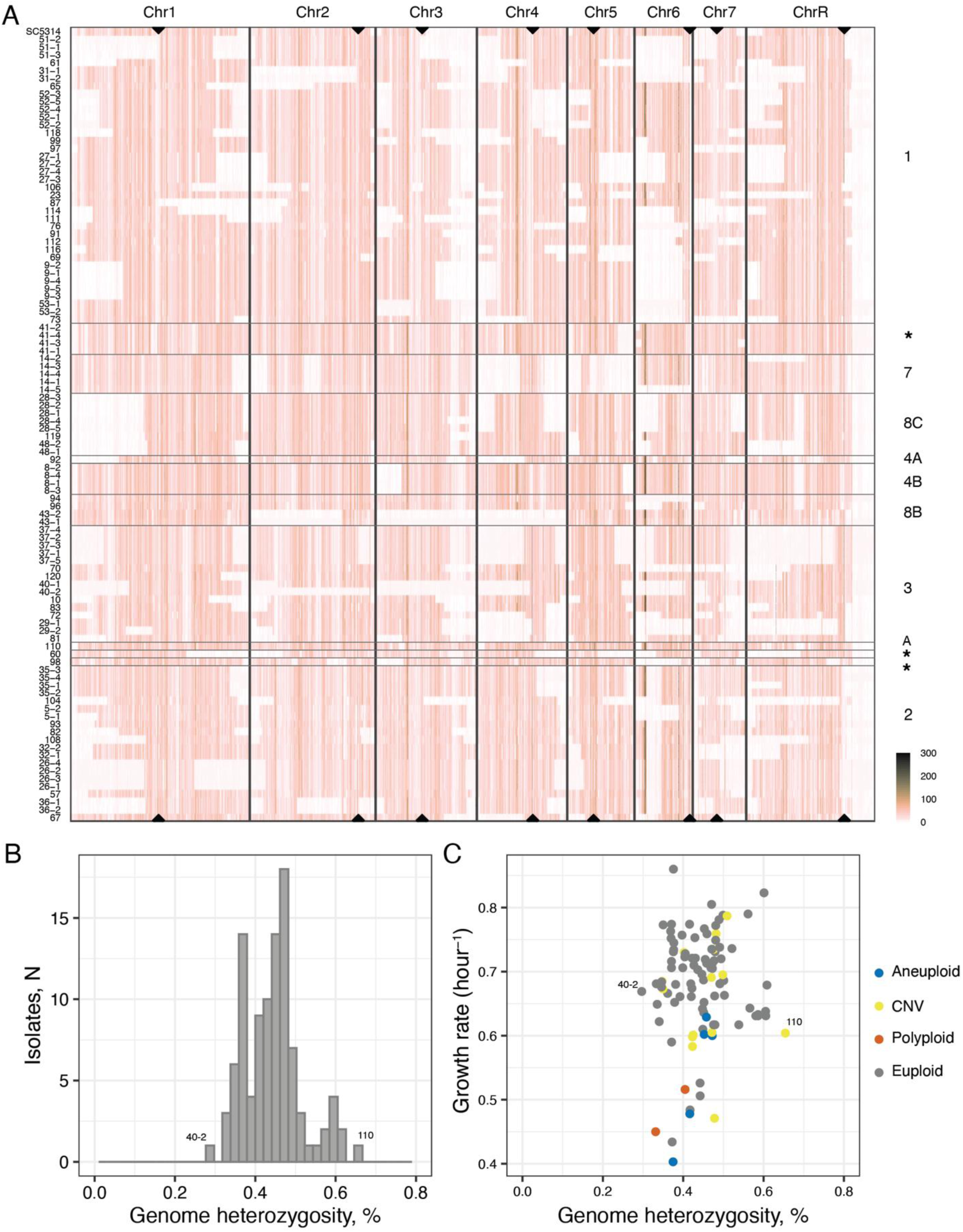
LOH patterns are often cluster specific. Genome heterozygosity values range from 0.3 to 0.65% and do not correlate with growth rate in rich media. (A) Genome-wide heterozygous SNP density for all isolates in 5 kb windows. Chromosome position is on the x-axis and isolates are in rows on the y-axis, ordered by the phylogenetic tree in Figure 1. Singleton isolates that do not belong to a cluster are designated with an asterisk. Isolate IDs are indicated on the left and cluster labels on the right of the figure. Centromeres are designated by black triangles on the top and bottom rows. SNP density is shaded from orange to dark brown. (B) Histogram of percent genome heterozygosity (see Methods and Supplemental Table S1). (C) Scatterplot of genome heterozygosity (x-axis) and growth rate/hour (y-axis). Isolates are colored by chromosome copy number. Isolates with the lowest (40-2) and highest (110) heterozygosity values are labeled.

All isolates were homozygous for the ChrR region located to the right of the rDNA locus. We identified multiple cluster-specific LOH regions, including a ∼450 kb segment of the far-right arm of Chr3 in cluster 1 and a ∼245 kb segment of the left arm of Chr6 in cluster 2. Clusters with small numbers of isolates also had large LOH blocks in all isolates (such as the left arm of Chr1 in cluster 8C), but this might be due to limited sampling rather than a cluster-specific LOH pattern. Finally, while serial isolates shared unique LOH fingerprints relative to the other isolates, forty-four percent of serial isolates contained at least one isolate that underwent LOH of at least one chromosome segment, highlighting the within-host heterogeneity of infecting populations.

### Genome-wide heterozygosity does not correlate with fitness

Higher genome-wide heterozygosity has been associated with fitness and virulence in *C. albicans* (8,24). For each isolate, we estimated the degree of genome-wide heterozygosity as the percentage of nucleotide positions having an allelic ratio between 0.25 and 0.75 (see Methods and Supplemental Table S1). Genome-wide heterozygosity ranged from 0.3 to 0.65% heterozygous nucleotide positions with a mean of 0.45% (Figure 2B). Heterozygosity of individual chromosomes ranged from a minimum of 0.015% on Chr4 (isolates 43-1 and 43-2) to a maximum of 0.86% on Chr6 (isolate 43-2) (Supplemental Table S1). Chr6 had the widest range of heterozygosity (0.02 to 0.86%) and ChrR had the lowest maximum heterozygosity (maximum 0.53% of all nucleotide positions), which reflects the LOH patterns seen in Figure 2A. Similar to previous studies, all isolates were homozygous between the ribosomal DNA locus (rDNA) and the right telomere of ChrR, which contributes to the lower mean heterozygosity of ChrR.

To investigate whether the shared genomic features of phylogenetic clusters such as LOH patterns were associated with shared phenotypes in those clusters, we tested the correlation of cluster membership and two different phenotypes: fluconazole tolerance measured by supra-MIC growth (SMG) and growth rate in rich media. There was no correlation between phylogenetic clusters and SMG (*p* = 0.06) or growth rate (*p* = 0.87).

We then tested the association of genome-wide and chromosomal-level heterozygosity of all euploid isolates and growth phenotypes (Supplemental Table S3). There was no statistically significant correlation between genome-wide heterozygosity and either SMG (*p* = 0.993) or growth rate (*p* = 0.48, see Figure 2C). However, at the individual chromosome level, fluconazole tolerance (SMG) was negatively correlated with heterozygosity of Chr1 (Pearson’s correlation coefficient = -0.25, *p* = 0.033) and Chr3 (Pearson’s correlation coefficient = -0.28, *p* = 0.016), but positively correlated with Chr4 heterozygosity (Pearson’s correlation coefficient = 0.35, *p* = 0.003).

### Heterozygous nonsense mutation of *ERG251* on Chr4 leads to high azole tolerance

Identifying mechanistic underpinnings of diverse *C. albicans* genotypes and specific phenotypes remains challenging. Isolate 120 had the highest fluconazole tolerance of any isolate as well as the second highest Chr4 heterozygosity (Figure 3A). We searched for genetic variants that could explain this isolate’s high tolerance. Isolate 120 was euploid and had 16 unique, high-impact single nucleotide variants (SNVs) compared to all 100 other clinical isolates in this study. One high-impact variant on Chr4 resulted in a heterozygous nonsense mutation in *ERG251* (W145*), encoding the C-4 sterol methyl oxidase important for ergosterol biosynthesis. We recently reported that heterozygous mutations in *ERG251* recurrently evolved during adaptation to fluconazole *in vitro*, and single allele dysfunction of *ERG251* caused multi-azole tolerance across diverse genetic backgrounds of *C. albicans* (37). To determine the extent to which the heterozygous nonsense mutation in *ERG251* was contributing to the azole tolerance phenotype, we complemented the nonsense allele with the wild-type *ERG251* allele in isolate 120. Two independent complemented strains had significantly decreased multi-azole tolerance compared to isolate 120 (Figure 3B one-way ANOVA *p* < 0.01, and Supplemental Figure S3). The heterozygous nonsense variant accounted for 60% of the tolerance phenotype (Figure 3B) with no significant fitness defect in the absence of drug (Supplemental Figure S3, one-way ANOVA, p = 0.658).

**Figure 3.**
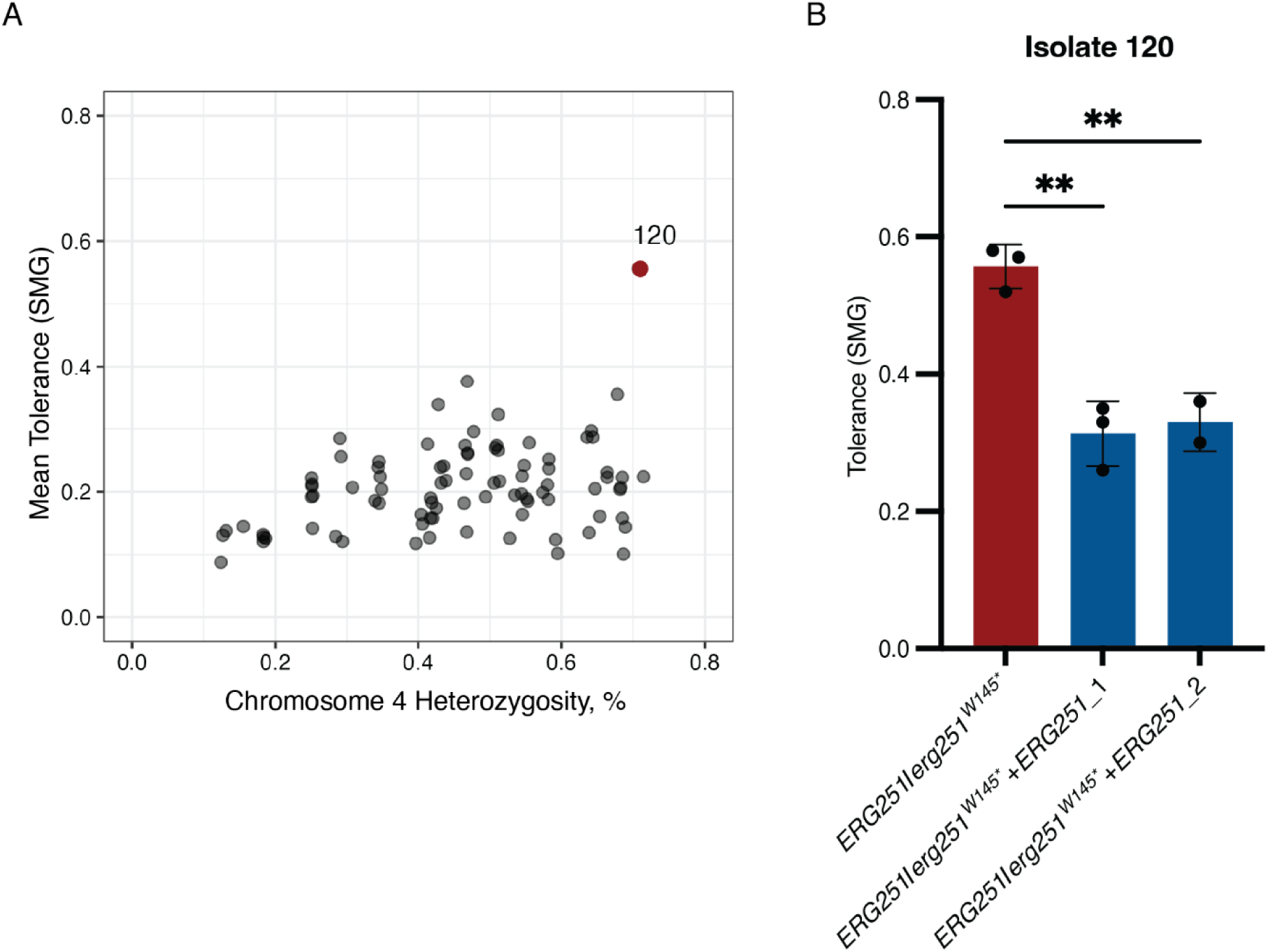
Isolate with *ERG251* heterozygous variant has highest fluconazole tolerance; wild-type *ERG251* complementation partially restores susceptibility. (A) Scatterplot of Chr4 heterozygosity % (x-axis) and mean 48-hour fluconazole SMG (y-axis). Isolate 120 has the highest fluconazole SMG (annotated). (B) 48-hour SMG of isolate 120 (*ERG251/erg251^W145*^*) and two wild-type complement strains (*ERG251/erg251^W145*^+ERG251*_1 and _2, two independent transformants). The SMG is the mean + SEM of at least two technical replicates per strain (represented as dots). One-way ANOVA with Dunnett’s multiple comparisons test was used to test for differences between isolate 120 *ERG251/erg251^W145*^* and the complement strains. ** *p* < 0.01.

### Karyotypic differences are common and difficult to detect using short-read sequencing data

To examine the structural genomic differences within the *C. albicans* bloodstream isolates, we performed karyotype analysis by pulsed field gel analysis. We observed between 7 and 11 electrophoretic bands in each isolate, ranging in size from ∼0.34 Mb to ∼3.19 Mb (Figure 4A). Mini chromosomes (bands smaller than SC5314 wild-type Chr7) ranging from ∼0.34 to ∼0.85 Mb in size were detected in seven isolates, including isolates 60 and 72, and all serial isolates isolated from case 35 (Figure 4A, red arrows). We analyzed WGS data for potential copy number changes (Figure 4B) or structural variants related to changes in electrophoretic band sizes. Using short-read paired-end Illumina sequencing data, high confidence recombination breakpoints could not successfully explain karyotypic differences observed by pulsed field electrophoresis (see Methods). For example, in isolate 60 a novel reciprocal recombination event between chromosomes 2 and 6 was detected from sequencing data (supplemental Figure S4), but this predicted recombination event did not support the banding pattern observed in Figure 4A.

**Figure 4.**
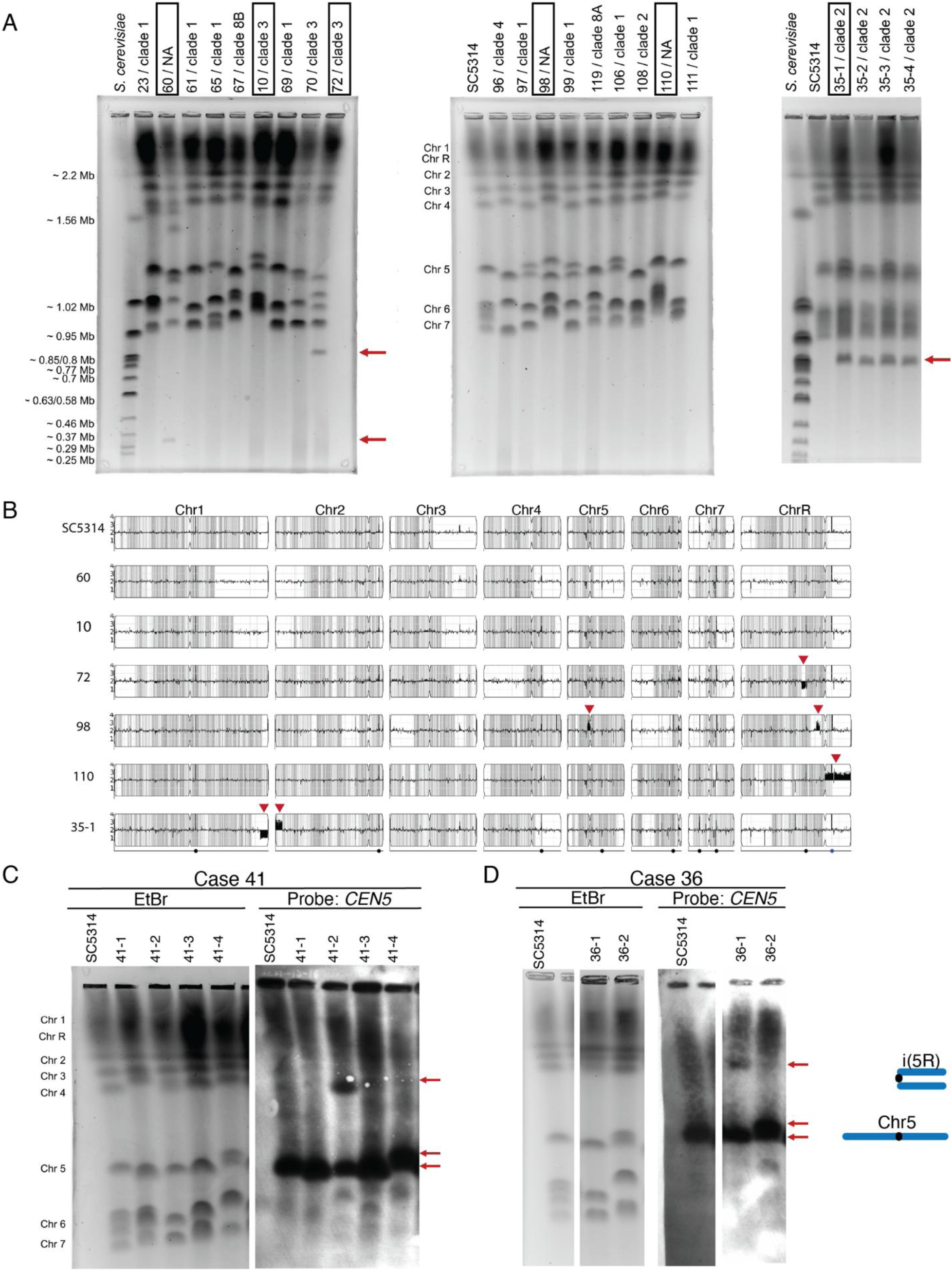
Karyotypic variation between serial isolates is not easily detected by short-read WGS analysis. (A) Three representative karyotype gels of clinical isolates stained with ethidium bromide. Novel mini chromosomes are indicated with red arrows in gels 1 (∼0.35 and ∼0.85 Mb bands) and 3 (∼0.84 Mb band). Isolate and cluster IDs are at the top of each lane. Reference genome karyotypes for *S. cerevisiae* (S288C genetic background) and *C. albicans* SC5314 are indicated with approximate sizes in Mb. (B) WGS data of representative isolates (IDs in boxes in A) plotted using YMAP. Relative copy number (y-axis) and heterozygous SNP density per 5 kb region in gray (x-axis). Karyotype gel and Southern blot analysis of (C) Case 41 and (D) Case 36. In all isolates, the *CEN5* probe hybridizes to 1 or 2 bands corresponding to the Chr5 homologs (lower red arrows). In isolate 41-2 and 36-1, the *CEN5* probe also hybridizes to a novel ∼1.6 Mb band (upper red arrow) that corresponds to the molecular weight of isochromosome 5R (i(5R), see schematic).

We examined the karyotypic patterns of all serial isolate cases and identified differences between isolates in 10 out of 18 cases. Representative serial isolate cases 41 and 36 have variation in bands corresponding to WT Chrs 5, 6 and 7 (Figure 4C and D). In both cases, one isolate acquired rearrangements involving the centromere of Chr5 (*CEN5*), as detected by Southern blotting (Figure 4C). All isolates had a single band corresponding to wild-type Chr5, but isolates 36-1 and 41-2 also had a larger ∼1.5 Mb band that hybridized to the *CEN5* probe (Figure 4C and 4D, upper red arrows). The ∼1.5 Mb band might represent a novel form of Chr5 – for example, an isochromosome consisting of two right arms of Chr5 (i(5R)) is ∼1.6 Mb (13). It might also be a more complicated translocation between multiple chromosomes, however neither scenario was confidently supported by short-read data.

### Serial isolates have differing aneuploidies and LOH patterns

The frequency of aneuploidy reported in clinical isolates has varied between studies. To identify potential aneuploidies, we analyzed the sequencing read depth of all isolates. Eight of 101 isolates were aneuploid for an entire chromosome, and 7 of these isolates were aneuploid for multiple chromosomes (Supplemental Table S4). Chromosomes 1, 5 and 7 were most commonly amplified.

Studies of aneuploidy in serial clinical isolates are uncommon (13,23,41), limiting our understanding of the dynamics of within-host aneuploidy during infection. Two serial isolate cases in this study had whole chromosome aneuploidy. Case 43 consisted of two isolates that were collected 17 days apart. Both isolates had Chr1 trisomy but differing copy number amplifications ranging from trisomy to tetrasomy of Chr5, 6 and 7, suggesting that aneuploidy was persistent in the patient for more than two weeks (Figure 5A).

**Figure 5.**
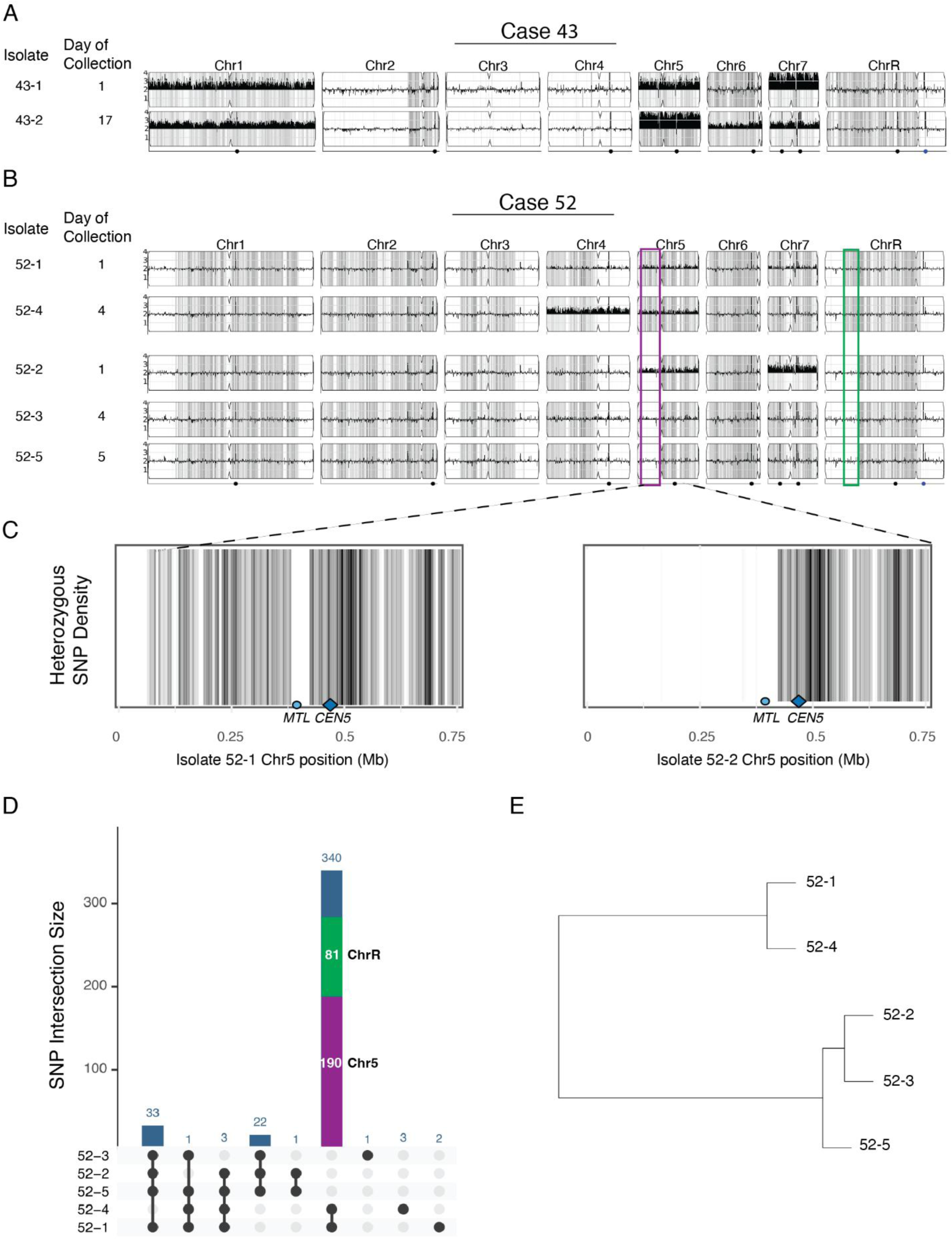
Aneuploidy and LOH differ between serial isolates collected within days from the same patient. (A and B) WGS data plotted with YMAP. Relative copy number (y-axis) and heterozygous SNP density per 5 kb region in gray (x-axis). Isolate ID and day of collection are indicated. (A) Case 43. (B) Case 52, ordered by genotype. Differing LOH patterns on Chr5 and on ChrR are outlined in purple and green respectively. (C) Chr5 *MTL* is involved in different LOH patterns. Heterozygous SNP density per 5 kb region in gray (x-axis) for isolates 52-1 and 52-2. (D) Shared SNP patterns distinguish sublineages within case 52. Upset plot of shared SNPs, where the bar plot is the number of positions shared among isolates connected by the dots below. Heterozygous and homozygous alternate alleles are counted together. SNPs shared by 52-1 and 52-4 corresponding to the LOH regions on Chr5 (purple) and ChrR (green) are indicated. (E) Dendrogram of case 52 isolates based on shared SNP positions.

Case 52 included 5 isolates collected over a 5-day period. Four of the isolates had Chr5 copy number than varied between 2 and 3 copies, suggesting instability of the aneuploidy either in the infecting population or during subculture (Figure 5B). Additionally, isolate 52-2 had Chr7 trisomy and isolate 52-4 had Chr4 trisomy (Figure 5B). The final isolate of the series, 52-5, was euploid for all chromosomes. In addition to aneuploidy differences, case 52 isolates had differences in LOH patterns and SNPs. Interestingly, all of case 52 isolates were homozygous for the *MTLa* allele on Chr5, but we identified two different LOH patterns involving the *MTL* locus (Figure 5C). Isolates 52-1 and 52-4 had a 42 kb LOH region of Chr5 which encompassed the ∼9 kb *MTL* locus (from approximately position Chr5:385200 to Chr5:427200). Isolates 52-2, 52-3, and 52-5 had a 430 kb LOH extending from the same breakpoint (Chr5:427200) and extending to the Chr5 left telomere. Isolates 52-2, 52-3 and 52-5 also shared an LOH event of the left arm of ChrR that was not present in 52-1 and 52-4. We quantified the SNPs present in fewer than all five isolates (unfixed SNPs) and determined their genetic relationship. The 5 isolates differed by only 406 SNPs and only 7 of the SNP differences were due to *de novo* mutations, with the remaining differences due to LOH (Figure 5D). Using shared SNP data, we generated a dendrogram for the isolates that demonstrates the existence of two different subpopulations sampled over the 5-day collection period of this bloodstream infection (Figure 5E).

### Breakpoints of CNVs are associated with repeat regions

Repetitive sequences are frequently associated with segmental chromosome copy number variants (CNVs) in laboratory-evolved *C. albicans* strains (15,16), so we determined the distance from each CNV breakpoint to the nearest repetitive region (see Methods). Sixteen isolates from 11 patient cases in this study had CNVs ranging in size from 34.6 kb to 1.13 Mb and involving chromosomes 1, 2, 5, 6, 7 and R (Supplemental Table S5). All CNV breakpoints were less than 35 kb from a repeat sequence (Supplemental Table S5). In 13 isolates, there were repetitive regions within 9.1 kb of all CNV breakpoints (Figure 6A). These clinical isolates support the association between CNVs and repetitive sequences in *C. albicans*.

**Figure 6.**
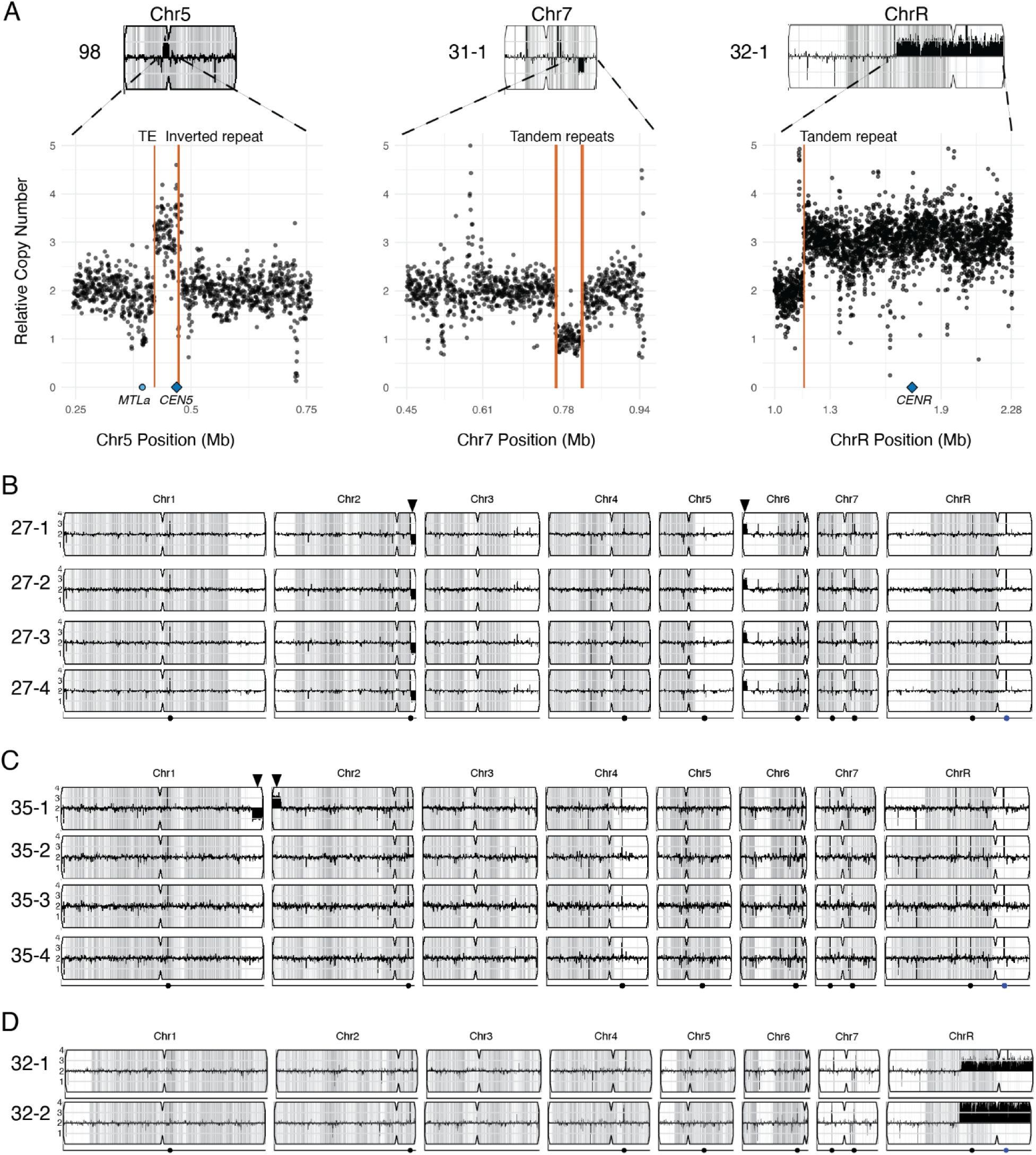
CNVs are associated with repeat regions and vary between serial isolates. (A - D) WGS data of serial isolate cases plotted using YMAP. Relative copy number (y-axis) and heterozygous SNP density per 5 kb region plotted in gray (x-axis). (A) Representative CNVs with nearest repeat regions indicated: Isolate 98 had a 60 kb amplification on Chr5 flanked by a transposable element (TE) and *CEN5* inverted repeat; Isolate 31-1 had a 54 kb deletion on Chr7 flanked on each side by tandem repeats; and Isolate 32-1 had a 1.13 Mb amplification on ChrR that begins on the left arm at a tandem repeat and extends through the centromere to the right telomere. (B) All serial isolates of case 27 had a 75 kb deletion of Chr2 and a 70 kb amplification of Chr6. (C) Only the first serial isolate of case 35 had a 165 kb deletion of Chr1 and a 148 kb amplification of Chr2. (D) A segmental amplification beginning in the left arm and extending to the right telomere of ChrR differs in amplitude between serial isolates 32-1 and 32-2.

### Copy number variation differs between serial isolates from the same patient

We further analyzed CNVs across serial and recurrent isolates collected from individual patients. Four patient cases had at least one serial isolate with a CNV. In case 27, all four isolates shared a 75 kb heterozygous segmental deletion that extended from the major repeat sequence (MRS) of Chr2 to the right telomere and a 70 kb amplification (3 copies) that extended from the left telomere of Chr6 (Figure 6B). In case 35 (4 total isolates), only isolate 35-1 had two large subtelomeric CNVs: Chr1 had a 160 kb heterozygous segmental deletion that extended from the right telomere, and Chr2 had a 140 kb segmental trisomy that extended from the left telomere; neither CNV was present in the other three serial isolates (Figure 6C). Case 32 had two isolates that were collected on the same day; both had amplification of the right arm of ChrR, with identical breakpoints but differing copy numbers (3 or 4 copies) of the amplified segment (Figure 6D). These results underscore that within-patient differences in both number and amplitude of CNVs are common.

### Individual gene amplifications and deletions are usually shared by serial isolates

Few genome-wide surveys of individual gene amplification or deletion events in *C. albicans* isolates exist (24). We analyzed read depth to identify copy number alterations spanning ≤ 10 kb and affecting 1 – 5 genes, focusing specifically on coding genes and excluding dubious ORFs. Ninety-four isolates had individual gene amplifications or deletions, with 1 to 5 genes affected per isolate (Supplemental Table S6). Amplifications were present in 7 different coding genes including *CYP5* (orf19.7421), predicted oxidoreductases orf19.113 and orf19.3131, and predicted zinc transporter orf19.3132, while the remaining three genes had unknown function (Supplemental Table S6). The most common gene amplification was *CYP5,* found in 41 of 101 isolates that were distributed across seven phylogenetic clusters.

There were deletions in nine different coding genes. Of the nine deleted genes, orf19.5736 is an ALS family adhesin, orf19.99 has predicted nucleotidase function, and the other seven have unknown function. The most commonly deleted gene was orf19.4069, a protein with unknown function, which was absent in 70 of 101 isolates (Supplemental Figure S5). Orf19.4069 was deleted in 95% of cluster 1 isolates, all cluster 2 and cluster 3 isolates, and two singleton isolates, suggesting there is cluster specificity in the distribution of this gene. We did not identify any instances of a gene being deleted in one isolate but duplicated in any other isolate.

We also quantified differences in gene-level amplifications/deletions between serial or recurrent isolates collected from individual patients. Only 2 of 19 serial or recurrent isolate cases had differing gene deletions between related isolates. Orf19.3924 (uncharacterized protein containing a steryl acetyl hydrolase domain) was absent in isolate 5-1 but present in isolate 5-2, collected on the same day. Orf19.5736 (*ALS5*) was present in isolates 14-1 through 14-4, but absent in isolate 14-5, collected from the same patient 338 days later. These results highlight the heterogeneity of gene content within and across *C. albicans* strains.

### Polyploidy in a recurrent infection is associated with a significant fitness cost

We identified whole genome duplication events in two isolates by DNA staining followed by flow cytometry. Both isolates were polyploid and highly aneuploid (Figures 7A and B): Isolate 76 was greater than tetraploid and Isolate 14-5 was triploid by flow cytometry. Notably, the triploid isolate 14-5 was collected 338 days after four serial isolates were first collected from this patient during the study period (Figure 7C). Isolates 14-1 through 14-4 were euploid diploid as determined by WGS read depth and flow cytometry analysis.

**Figure 7.**
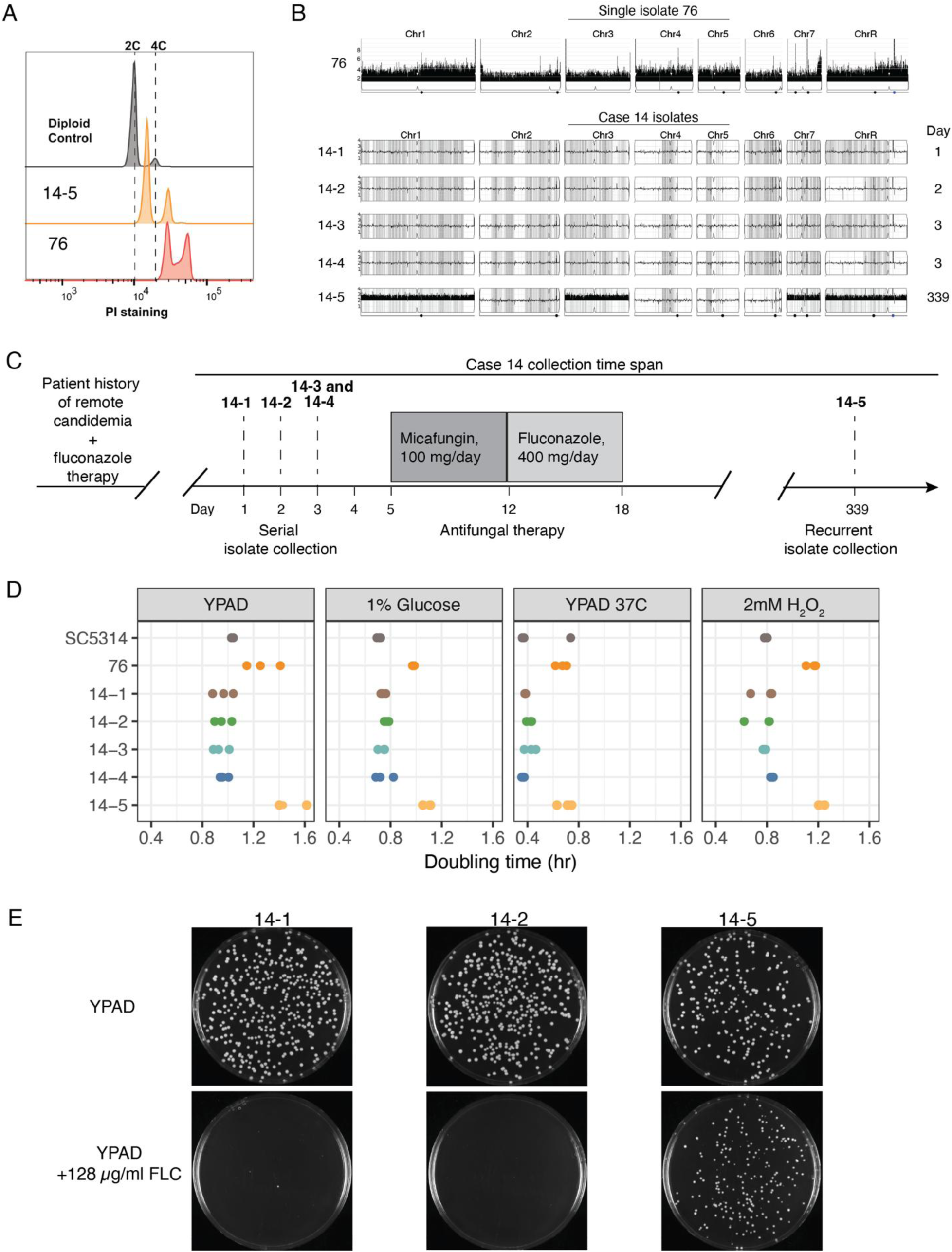
Within-host polyploidy, including polyploidy arising within a recurrent infection. (A) Ploidy analysis by flow cytometry with the diploid control and polypoid isolates 14-5 and 76. (B) WGS data of cases 76 and 14 plotted using YMAP. Relative copy number (y-axis) and SNP density per 5 kb region (x-axis). Isolate 76 was greater than tetraploid with chromosome relative copy numbers ranging from 3.4 to 4.7 (Supplemental Table S4). Isolate 14-5 was near triploid with chromosome copy numbers ranging from 2.1 to 3.3 (Supplemental Table S4). (C) Patient history, candidemia isolate collection and antifungal therapy for case 14. The patient had a remote history of candidemia treated by fluconazole (FLC) before the current study began. Isolates 14-1 through 14-4 were collected prior to repeat micafungin and fluconazole therapy and isolate 14-5 was collected 338 days later. (D) Doubling time in hours (x-axis) of isolates grown in rich media (YPAD), 1% glucose, 2mM H_2_0_2_, and 37°C. (E) Growth of FLC-susceptible diploid (14-1, 14-2) and polyploid (14-5) isolates on YPAD at 48 hours, YPAD + 8 μg/mL FLC at 72 hours, and YPAD + 128 μg/mL FLC at 72 hours.

To investigate the genetic relatedness over the collection timeframe, we quantified SNP differences between the isolates. Approximately 2,800 total SNPs separated the five isolates in case 14 and more than 2,700 of these were due to only two LOH events. Specifically, isolate 14-5 had an LOH event involving most of Chr6 relative to the other 4 isolates, and isolates 14-2 and 14-4 shared an LOH event of the left arm of ChrR (Figure 7B). There were fewer than 50 *de novo* SNPs unique to any single isolate from this patient, despite the intervening antifungal therapy and hundreds of days separating isolate 14-5 from the earlier serial isolates. All five isolates were sensitive to fluconazole, micafungin and amphotericin B, and all isolates had a fluconazole SMG of <0.15 (Supplemental Table S1), indicating that isolate 14-5 had no *in vitro* fitness advantage in the presence of antifungal drug.

To determine the magnitude of fitness effects in both polyploid isolates, we quantified their growth relative to the reference strain SC5314 and to the euploid diploid isolates 14-1 through 14-4. Both polyploid isolates had significantly longer doubling times than all diploid isolates in rich media and in the presence of low glucose, oxidative stress, and thermal stress (Figure 7D, *p* < 0.01, two-way ANOVA with Tukey’s HSD post-hoc). Polyploid cells typically have increased genome instability relative to diploid cells, therefore we hypothesized that the polyploid isolates would generate more growth-promoting genetic diversity in the presence of antifungal drug stress than their related diploids. To test this hypothesis, we grew the drug susceptible diploid (14-1, 14-2) and polyploid (14-5) isolates overnight in YPAD and then plated them on agar plates containing YPAD, YPAD + 8 μg/mL fluconazole, or YPAD + 128 μg/mL fluconazole and quantified colony formation (Figure 7E and Supplemental Table S8). Polyploid isolate 14-5 gave rise to significantly more colonies than the diploid isolates 14-1 and 14-2 in the presence of low and high concentrations of fluconazole (p < 0.01, two-way ANOVA with Tukey’s HSD post-hoc). The proportion of colonies that grew on 128 μg/mL fluconazole relative to colonies that grew on YPAD alone was 78% for polyploid isolate 14-5, but only 0-2% for diploid isolates 14-1 and 14-2 (Supplemental Table S8). Our results indicate that a drug-sensitive polyploid isolate can rapidly generate progeny with phenotypic diversity, including a fitness advantage in presence of drug.

## Discussion

We used comparative genomic and phenotypic analyses to comprehensively evaluate 101 *C. albicans* bloodstream isolates from an academic medical system in Minnesota’s Twin Cities region. The extensive genetic diversity represented in our local isolates is consistent with a 2018 WGS analysis of 182 globally-collected *C. albicans* clinical, commensal and environmental isolates and a 2023 WGS analysis of 45 commensal isolates, highlighting the global spread of this opportunistic pathogen (6,8). Most of our study isolates clustered with several previously described clades of *C. albicans* and helped refine clades 4 and 8 into five separate clusters. While our study methods were limited to single colony bloodstream isolates, we identified 6 isolates that do not belong to any known phylogenetic clusters.

We determined genome-wide heterozygosity of all isolates by calculating the allele frequency of nucleotide positions across the entire genome. The heterozygosity values calculated in this study differ substantially from previous reports that used different methods (24), but are similar to genome characteristics estimated from the publicly available genome profiling tool GenomeScope (42). We found no association between genome-wide heterozygosity and growth rate, in contrast to Hirakawa et al. (2015), which might be due to the differences in methodology and/or sample size. We identified three significant correlations between chromosome-level heterozygosity and fluconazole tolerance, including a positive correlation with Chr4 and negative correlations with Chr1 and Chr3. The isolate with the highest fluconazole tolerance in our study had a heterozygous *ERG251* nonsense mutation, and complementation with wild-type *ERG251* indicated that the nonsense allele accounted for ∼60% of the tolerance phenotype. Additional variants in this isolate might further contribute to the tolerance phenotype. Single allele dysfunction of *ERG251* confers drug tolerance without significant fitness costs across diverse isolates of *C. albicans*, but homozygous *ERG251* loss of function results in impaired biofilm formation, decreased virulence and decreased fitness in the absence of drug, further suggesting that maintenance of heterozygosity at this gene is important for pathogenicity and drug tolerance in the host (37,43).

Through karyotypic analysis, we identified extensive chromosomal variation among isolates. More than half of the serial isolate cases had karyotypic differences between related isolates, suggesting that extensive within-host genomic rearrangements commonly occur. Similarly, we found that whole and segmental chromosome copy number changes vary between serial isolates, even over very short collection time periods. These differences in copy number between serial isolates could be due to the instability of aneuploidy during laboratory culture or could represent within-host competition of sublineages with different fitness effects (44,45). We found that CNV breakpoints are associated with long repeat sequences, and this supports our previous reports that recombination between long repeat sequences contributes to both copy number amplification and deletion events in *C. albicans* (15,16,20,46). Ultimately, long read sequencing is required to resolve recombination events at long repeat sequences (47) and we propose that a pangenome approach is greatly needed to capture the full spectrum of structural variation across the species.

We identified within-patient acquisition of polyploidy in a recurrent infection occurring 338 days after initial isolates were collected. The polyploid isolate differed by only 41 *de novo* SNPs from the initial isolates and was identified long after both 7-day micafungin and 6-day fluconazole therapies. No prior studies have genomic evidence supporting the evolution of polyploid cells during a persistent human infection, however fluconazole treatment in a murine model of systemic infection resulted in the evolution of polyploid cells from a diploid progenitor (22,48). Fluconazole, the recommended step-down therapy for candidemia, is known to induce cell cycle defects and polyploidization in *C. albicans*, which could be an explanation for the ploidy change seen in this recurrent infection (49–51). While the polyploid isolate had significant fitness costs in the absence of drug *in vitro*, it rapidly generated phenotypic diversity leading to drug-resistant progeny. Because both polyploid isolates in this study were aneuploid for multiple chromosomes it is challenging to dissect the fitness effects of polyploidy from aneuploidy *per se*, however multiple studies support that polyploidization followed by rapid chromosome reshuffling promotes somatic genomic diversity and can increase the speed of adaptation in *C. albicans* and other fungi (22,48,49,52–54).

To our knowledge, this is the first genomic survey of *C. albicans* clinical bloodstream isolates from Minnesota’s Twin Cities area hospitals. An important strength of our study is the number of serial isolates collected from individual patients, providing novel insights into the within-host variation and population heterogeneity of bloodstream infections. We identified large-scale genomic changes that differed between closely related serial isolates including polyploidy, aneuploidy, CNVs and LOH.

## Conclusion

Our comprehensive serial isolate analysis demonstrates the extensive variation that exists during invasive infection and receives only limited clinical attention. This genomic variation could allow for rapid adaptation to stress and potentially complicate treatment even in the absence of acquired antifungal drug resistance. The within-host genomic diversity revealed by our study exposes critical limitations in standard clinical sampling and antifungal susceptibility testing, underscoring the need for more frequent and dynamic susceptibility assessments during treatment of persistent candidemia.

## Methods

### Isolate collection

Isolates were collected and stored as glycerol stocks as previously described (IRB-reviewed study IDs STUDY0000647 and STUDY00021428) (38). Isolates are single-colony subcultures of individual blood culture samples.

### MIC assay

Cultures were grown on Sabaraud’s Dextrose Agar plates at 35℃ for 24-28 hours and diluted in dH2O to a final optical density (OD600) of 0.1. Cell suspensions were diluted 1:2 into a 96-well NUNC plate containing 2x RPMI media (Cytiva, product no. SH30011.02) with or without drug. Drug concentrations ranged from 0.5μg/ml to 256 μg/ml FLC (Alfa Aesar, product no. J62015). Cells were incubated in a humidified chamber at 35℃ for 48h. At 24 hours, cells were resuspended and OD530 readings were taken and repeated at 48 hours. Each isolate was assayed in triplicate. The MIC50 of each strain was determined as the drug concentration at which ≥ 50% of growth was inhibited relative to the no-drug control at 24 hours post-inoculation. Supra-MIC growth (SMG) was measured as the average growth for all wells above the MIC50 after normalization to the no-drug control at 48 hours post-inoculation.

### Growth curve analysis

Overnight cultures were grown in a shaking incubator at 30°C in liquid YPAD medium with 2% dextrose (10 g/L yeast extract, 20 g/L Bacto peptone, 20 g/L dextrose, 0.04 g/L adenine and 0.08 g/L uridine), diluted to an OD600 of 0.1 and inoculated into 96-well plates with or without chemical stressor. Chemical stressors include 1% YPAD + 2mM H2O2 at 30°C, 0.1% carbon YPAD at 30°C, and 1% YPAD at 37°C. Cells were incubated at 30℃ in a BioTek Epoch 2 microplate spectrophotometer shaking in a double orbital (237rpm) with OD600 readings taken every 15 minutes for 48 hrs. Each isolate was assayed in triplicate.

### Transformation: Complementation o*f C. albicans ERG251* WT allele

Complementation of isolate 120 (clinical isolate with a heterozygous *erg251* nonsense mutation) was performed as previously described (37). Primers are provided in Supplemental Table S7 (37). Briefly, the wild-type *ERG251* gene, upstream region plus nourseothricin resistance gene (*NAT)* and downstream region of *ERG251* were PCR amplified from pAS3118 using primers 1652+1653. The subsequent *ERG251-NAT* construct was transformed into isolate 120. NAT-resistant transformants were PCR screened for correct integration of the *ERG251-NAT* construct using primer pairs 1634+1154 (left integration), 1636+1653 (right integration), and 1634+1653 (across integration). Transformants were validated by Sanger sequencing for correct integration.

### Karyotype analysis using Contour-Clamped Homogeneous Electric Field (CHEF)

Sample plug preparation was performed as previously (55). *C. albicans* SC5314 (lab ID AMS2401) and *S. cerevisiae* strain AMS2123 (S288C genetic background) were used as karyotype ladders. Overnight cultures were grown in a shaking incubator at 30°C to stationary phase, suspended in 1.5% low-melt agarose (Bio-Rad, product no. 161-3111), digested with 50 mg/mL Zymolyase (US Biological, product no. Z10001G) for 16 hours at 37°C, washed twice with 50mM ETDA (Fisher Scientific, product no. BP2482-1), incubated with 0.2 mg/mL proteinase K (Fisher Scientific, product no. BP1700100) and washed with 50mM EDTA before storage at 4°C. Whole chromosomes were separated in a 1% megabase agarose gel (Bio-Rad, product no. 1613109) in 0.5X TBE (Fisher Scientific, product nos. BP152-1, BP2482-1, 327130010) using a Bio-Rad CHEF-DRIII Pulsed Field Electrophoresis System (55,56). Run settings were Block 1: 60–120 36 hours 6V/cm 120° and Block 2: 120–300 12 hours 4.5 V/cm 120°. After 48 hours gels were stained with ethidium bromide (Invitrogen, product no. 15585-011) and imaged on a Bio-Rad Gel Doc XR+ System using a UV transilluminator light. Image colors have been inverted and saturation levels adjusted for clarity.

### Southern blot hybridization

DNA was transferred from CHEF gels to a BrightStar Plus nylon membrane (Invitrogen, product no. AM10102). Probe hybridization and detection were performed as previously described (15,16,41,55). Probes were constructed by PCR using DIG-11-dUTP nucleotides (Roche PCR Dig Probe Synthesis Kit, product no. 11636090910) and *CEN5* primer pair (forward: CGGTCCCTACGTTCGTCAAA, reverse: ACAGCCTCGTTGACCAGAAG) according to the manufacturer’s instructions. Southern blot film images were captured on a Bio-Rad Gel Doc XR+ System and saturation levels were adjusted for clarity.

### Ploidy determination by flow cytometry analysis

Cells were prepared as described previously (57). Overnight cultures were grown in a shaking incubator at 30°C to a cell density of 1×10^7^ cells/mL. Cultures were gently spun down at 3000 rpm for 3 minutes and the supernatant was removed. Cell pellets were resuspended in 70% ethanol, then washed twice with 50 mM sodium citrate (Fisher Scientific, product no. BP327-1). Following washing, cells were spun down and resuspended in 50 mM sodium citrate containing 0.5 mg/ml RNase A (Invitrogen, product no. 12091-021). Cells were treated with RNase A at 37°C for at least 2 hrs, then stained with 25 µg/ml propidium iodide (PI) (Invitrogen, product no. P3566) at 37°C in the dark overnight. Samples were diluted in 50 mM sodium citrate, and at least 10,000 singlets were analyzed using a Cytek Aurora flow cytometer (R0021). The 488-nm laser was used to excite the PI and 618/24 filters were used to detect the PI emission. Unstained cells were used as the negative control. Wild-type diploid SC5314 and haploid YJB12814 (58) were used as ploidy controls. Data were analyzed using FlowJo (https://www.flowjo.com/solutions/flowjo/downloads) (v10.8.1).

### DNA extraction and Illumina whole-genome sequencing

Overnight cultures were grown in a shaking incubator at 30°C. Genomic DNA was isolated using phenol-chloroform extraction as previously described (59). Libraries were prepared using the Illumina DNA Prep kit and Integrated DNA Technologies (IDT) 10-bp unique dual indexes (UDIs). Samples were sequenced on an Illumina NextSeq 2000 to produce paired-end 151-bp reads. Bcl-convert (v3.9.3) was used for demultiplexing, quality control and adapter trimming (60). Adapter and quality trimming were performed with BBDuk (v38.39) according to BBTools data preprocessing guidelines (61). Trimmed reads were aligned to the *C. albicans* SC5314 reference genome (SC5314_version_A21-s02-m09-r08) using BWA-MEM (v0.7.17) with default parameters (39,62). Aligned reads were sorted, duplicate reads were marked, and BAM output files were indexed with Samtools (v1.10) (63). Summary statistics were generated with Samtools, FastQC (v0.11.7) and Qualimap (v2.2.2-dev) and aggregated into a report using MultiQC (v1.16) (63–66). The average WGS coverage was 86x (range of 33x to 130x, Supplemental Table S1).

### CFU counting and colony morphology

Tetraploid isolate 14-5 and diploid isolates 14-1 and 14-2 were inoculated in YPAD from frozen stock and incubated for 36hr at 30°C and 220 rpm. Cultures were diluted in PBS to an OD_600_ of 0.01 followed by 1:100 dilution. A volume of 100μl of this dilution was plated on YPAD, YPAD+8μg/ml FLC, and YPAD+128μg/ml FLC agar plates. Three replicates were performed. Plates were imaged after incubation at 30°C for 48hr and 72hr using a BioRad GelDoc XR+ imaging system.

### Public data sets

Two hundred two additional publicly available *C. albicans* genomes from BioProjects PRJNA193498 (WGS SRA runs with library name “Pond-NNN”) and PRJNA432884 (all available SRA runs) were downloaded from NCBI Sequence Read Archive using the SRA toolkit (v3.0.0) (8,24,67). Sequencing data were processed as describe above and samples with < 20x mean coverage were discarded.

### Small variant calling and multilocus sequence typing

SNPs and small indels were called using GATK (v4.4.0) (68). SNPs were filtered on the parameters QD < 2.00, QUAL < 30.0, SOR > 3.0, FS > 60.0, MQ < 40.0, MQRankSum < -12.5, and ReadPosRankSum < -8.0. Indels were filtered on the parameters QD < 2.0, QUAL < 30.0, FS > 200.0, and ReadPosRankSum < -20.0. Bcftools (v1.17) was used to calculate variant allele frequency (VAF) per sample, filter for heterozygous variants with VAF between 0.15 and 0.85 and homozygous variants with VAF > 0.98, and to exclude known repetitive regions as annotated in the SC5314 A21-s02-m09-r08 GFF (rRNA, repeat_region, retrotransposon) and telomere-proximal regions, defined here as extending from each chromosome end to the first non-repetitive genome feature (15,69,70). The filtered variants were annotated using SnpEff (v5.0e, database built manually from SC5314 version A21-s02-m09-r09 with alternate yeast nuclear codon table (71).

The MLST loci for *C. albicans* are *AAT1* (orf19.11037), *ACC1* (orf19.7466), *ADP1* (orf19.459), *PMI1* (orf19.1390), *ALA1* (orf19.5746), *VPS13* (orf19.4416), and *ZWF1* (orf19.4752) (40,70). For each isolate, a consensus sequence was generated using bcftools and the PubMLST typing database was queried to find allelic matches for each locus and determine ST profiles. New alleles and MLST profiles were submitted to PubMLST. Isolate records were submitted to PubMLST and can be found in the isolate collection database (isolate field = study code from Supplemental Table S1).

### Phylogenetic tree construction, cluster determination and visualization

The filtered VCF file was subset to include only SNP sites with no missing data and converted to PHYLIP format using vcf2phylip (v2.8) (72). A maximum-likelihood tree was built using RAxML (v8.2.11) with the GTR+GAMMA model of nucleotide substitution and rapid bootstrapping (100x) (73). No outgroup was specified.

To define clusters, we adapted the approach described by Schikora-Tamarit and Gabaldón (2024). We defined cluster nodes as internal nodes with bootstrap support >95%, having subtending branch lengths above a minimum threshold, and having no child nodes that also met the definition for cluster nodes. We compared the results of using branch length and the ‘relative_branch_length’ defined by Schikora-Tamarit and Gabaldón (2024), which is branch length normalized to the longest distance between any two nodes of the tree. Since results agreed between actual branch length and relative branch length and were dependent on manual selection of a minimum threshold scaled to each approach, we used actual branch length. We compared a range of minimum threshold values to identify a value (branch length > 0.057) that had the most agreement with existing MLST and WGS-based clusters (7,8) and minimized the number of singleton isolates. Midpoint rooting of the RAxML bipartition file was performed with the R package *phangorn* (v2.11.1) (74,75). Isolate metadata was added to create a treedata object which was visualized using the R packages *ggtree* (v0.4.6) and *tidytree* (v3.10.1) (76).

### Recombination breakpoint analysis using Illumina paired reads

To identify potential structural variants observed by CHEF, an analysis of Illumina read pairs was performed (77). Read pairs that mapped to different chromosomes were extracted from the alignment file using samtools (v1.17), excluding reads mapping equally to multiple positions. Recombination breakpoints were determined by extracting separated read pairs clustering within 15 bp of a breakpoint and requiring 10 supporting reads on either end of the breakpoint. Due to the limitations of short read data, breakpoints between two chromosomes occurring near repetitive elements were filtered out, including annotated repeats in the reference genome and transposable elements identified with RepeatMasker (v4.1.1), using the transposon library curated by Oggenfuss et al (47). The predicted size changes of each chromosome due to isolate-specific recombination relative to the reference were determined by identifying paths from each telomere to another telomere, allowing a path to cross interchromosomal breakpoints, where the resulting path lengths were summed. Predicted chromosome sizes were compared to the band sizes observed with CHEF electrophoresis.

### Relative copy number analysis

Genome-wide copy number was visualized using YMAP. Then, sequencing read depth was used to identify relative copy number (78). GC content bias was calculated and corrected using deepTools (v3.5.4). Read depth was computed using Samtools depth with option -aa to output all positions. The rolling mean of nuclear read depth was calculated over 500 bp tiled windows using the R package *RcppRoll* (v0.3.0) (79). The median depth was calculated for each chromosome, and for the entire nuclear genome. To minimize bias due to chromosome-level copy number changes, median nuclear depth was then filtered to include only chromosomes with a depth between 0.85 and 1.15 of the raw median nuclear value. Relative read depth for each window was calculated by dividing the window’s read depth by the filtered median nuclear depth.

For individual genes, candidate ORFs were filtered from the SC5314 A21 genome feature file (GFF) to exclude repetitive subtelomeric regions, annotated repeat regions and ORFs annotated as dubious or transposable elements (70). Thresholds for changes in relative depth were set as < 0.25 or > 1.5, excluding chromosomes with known copy number changes. Candidate ORFs that underwent copy number change across at least 75% of the ORF were determined over a sliding window of 500 bp using R packages *GenomicRanges* (v1.54.1) and *rtracklayer* (v1.62.0) (80,81). Candidate CNVs were visually inspected in IGV for at least one representative isolate (82).

### Allele frequency calculation and genome heterozygosity estimation

We first attempted to estimate heterozygosity using GenomeScope, a reference-free method that infers genome characteristics from k-mer profiles (42). However, several isolates did not generate clear bimodal distributions of k-mers as expected in a diploid when using this tool, regardless of parameter settings or sequencing coverage. Therefore, we estimated heterozygosity using the aligned reads for each isolate. Samtools pileup files were generated from BAM files, with option -q (minimum mapping quality) of 60 (63). Counts of all bases at each position were generated using a python script adapted from https://github.com/berman-lab/ymap/tree/master (83). Allele frequencies were calculated for each position. Minimum (25%) and maximum (75%) allele frequencies were set for counting diploid positions as heterozygous and scaled to account for ploidy changes, similar to limits implemented in Yeast Mapping Analysis Pipeline (YMAP) (83). Heterozygous positions were counted in 5,000 bp tiled windows for all isolates. Values were binned in 10% increments up to the maximum window value. The sum of heterozygous positions for each chromosome was divided by the size of the chromosome to calculate chromosome heterozygosity as a percentage. All chromosome heterozygous positions were summed and divided by the size of the nuclear genome (SC5314 A21 fasta) to calculate genome-wide heterozygosity as a percentage.

### Statistical analysis

Correlation testing was performed on phenotypic data collected from diploid isolates with no large CNVs (38). Pearson’s correlation coefficient was calculated for growth rate, doubling time and fluconazole SMG relative to genome heterozygosity and individual chromosome heterozygosity. Spearman’s correlation was calculated for growth rate and fluconazole SMG relative to clade membership. Multiple test correction was performed using the Holm method (84). Correlation analysis was performed with the R package *correlation* (85).

GraphPad Prism (v.10.11) for macOS was used to calculate differences between *ERG251* heterozygous variant clinical isolates and wild-type engineered strains using one-way ANOVA followed by Dunnet’s multiple comparisons test. The R package *rstatix* (v0.7.2) was used to calculate differences between polyploid and diploid isolates in multiple growth conditions using two-way ANOVA followed by Tukey HSD for pairwise comparisons (86).

## Declarations

## Ethics approval

Sample and data collection were approved by the University of Minnesota Institutional Review Board (study IDs STUDY0000647 and STUDY00021428).

## Consent for publication

Not applicable

## Availability of data and materials

The WGS data generated during this study has been deposited at the NCBI Sequence Read Archive under BioProject ID PRJNA1068683. Phenotypic data generated during this study are included in the published article and its supplementary files. Analysis scripts are available at https://github.com/selmeckilab/2024_Candida_albicans_MEC_analysis

## Author contributions

Conceptualization: NES, SE, SK, AS. Sample collection: SE, EW. Experiments: EW, XZ, CZ. Data analysis: NES, EW, XZ, CZ. Visualization: NES, XZ, EW. Writing – first draft: NES. Writing – review and editing: NES, XZ, EW, CZ, SE, SK, AS.

## Acknowledgements

We are grateful to members of the Selmecki laboratory for feedback on methodology, data presentation, and clarity. We thank Dalton Piotter for preparation and imaging of CHEF gels and Southern blots. The Minnesota Supercomputing Institute (MSI) at the University of Minnesota provided essential resources that contributed to the research results reported within this paper.

