## Supplemental figures S1-S6 for "Aneuploidy, polyploidy, and loss of heterozygosity distinguish serial bloodstream isolates of *Candida albicans*"

**Supplemental Figure S1 (Scott *et al.*)**

**Aneuploidy, polyploidy, and loss of heterozygosity distinguish serial bloodstream isolates of *Candida albicans***

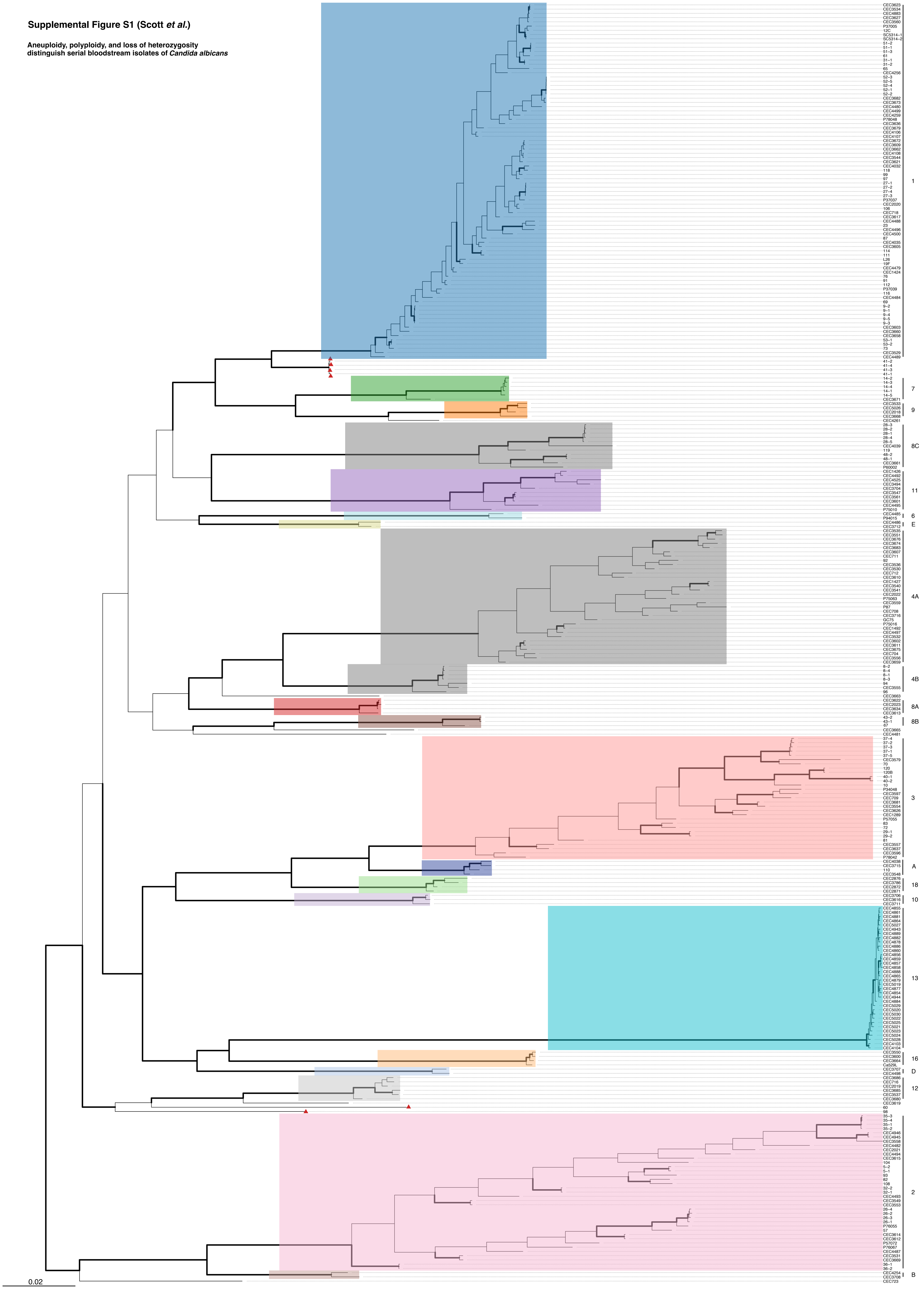


**Supplemental Figure S1**. Maximum-likelihood, midpoint-rooted phylogenetic tree with 101 new isolates (this study) and 199 publicly available genomes (Supplemental Table S2). Cluster labels correspond to Odds et al. (2007) and Ropars et al. (2018). Six isolates (red triangles) from three cases do not cluster with any previously described clade (patient cases 41, 60 and 98). MLST clade 4 is divided into 2 clusters designated 4A and 4B by this analysis. MLST clade 8 is divided into 3 clusters designated 8A, 8B and 8C. Bootstrap support > 95% represented by thick, bolded bars. The scale bar represents the number of substitutions per site.

Supplemental Figure S2 (1 of 7)

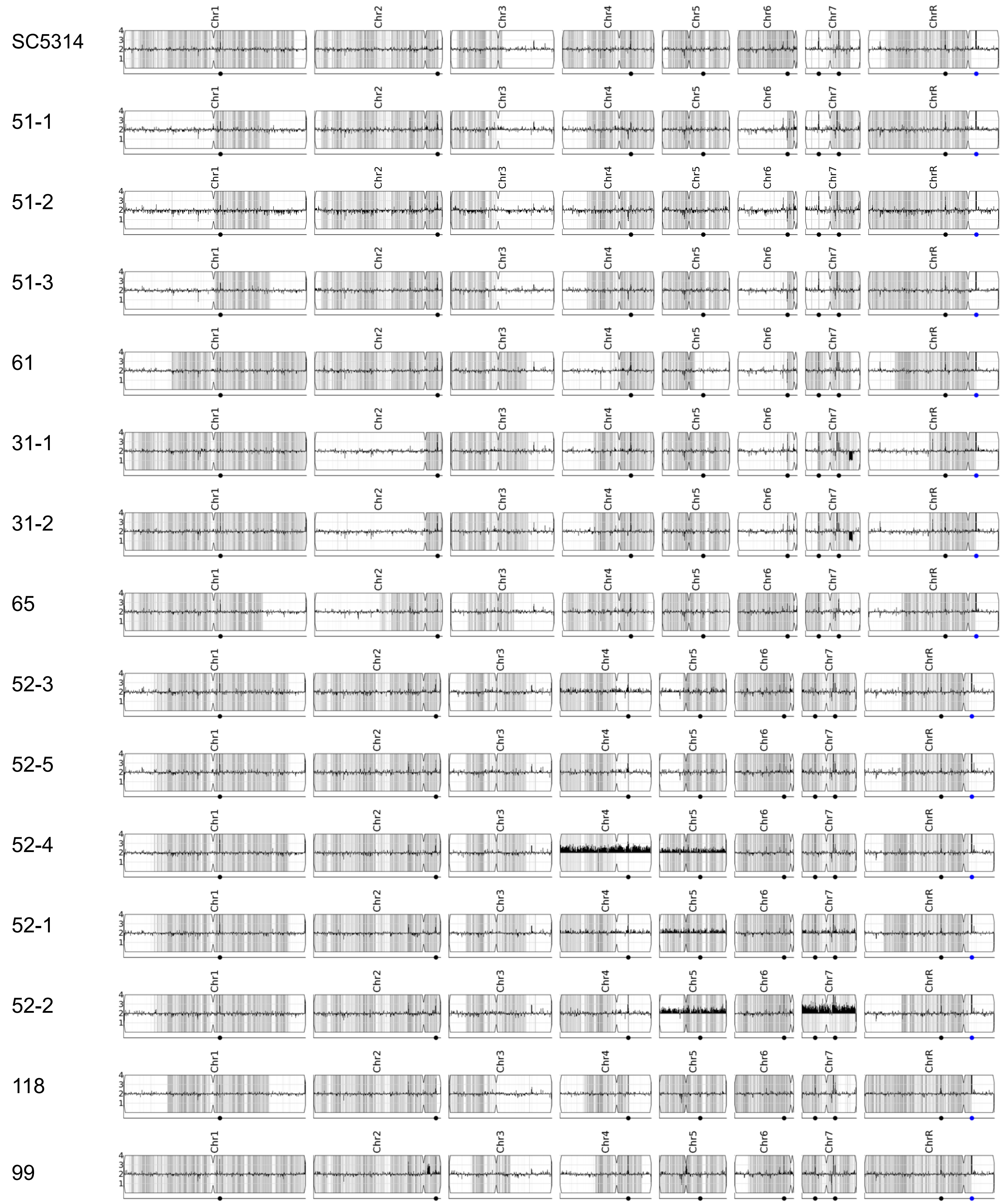

Supplemental Figure S2 (2 of 7)

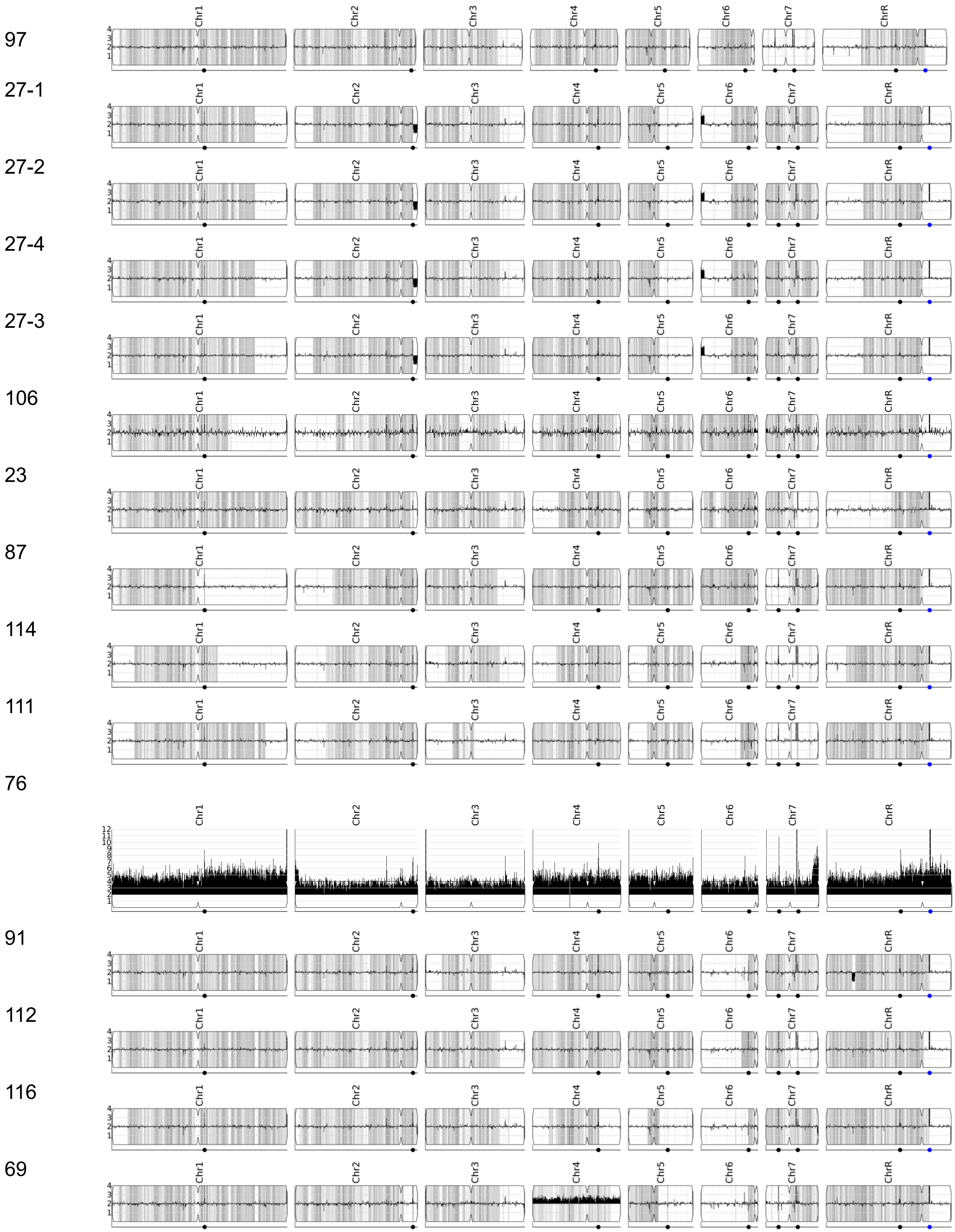

Supplemental Figure S2 (3 of 7)

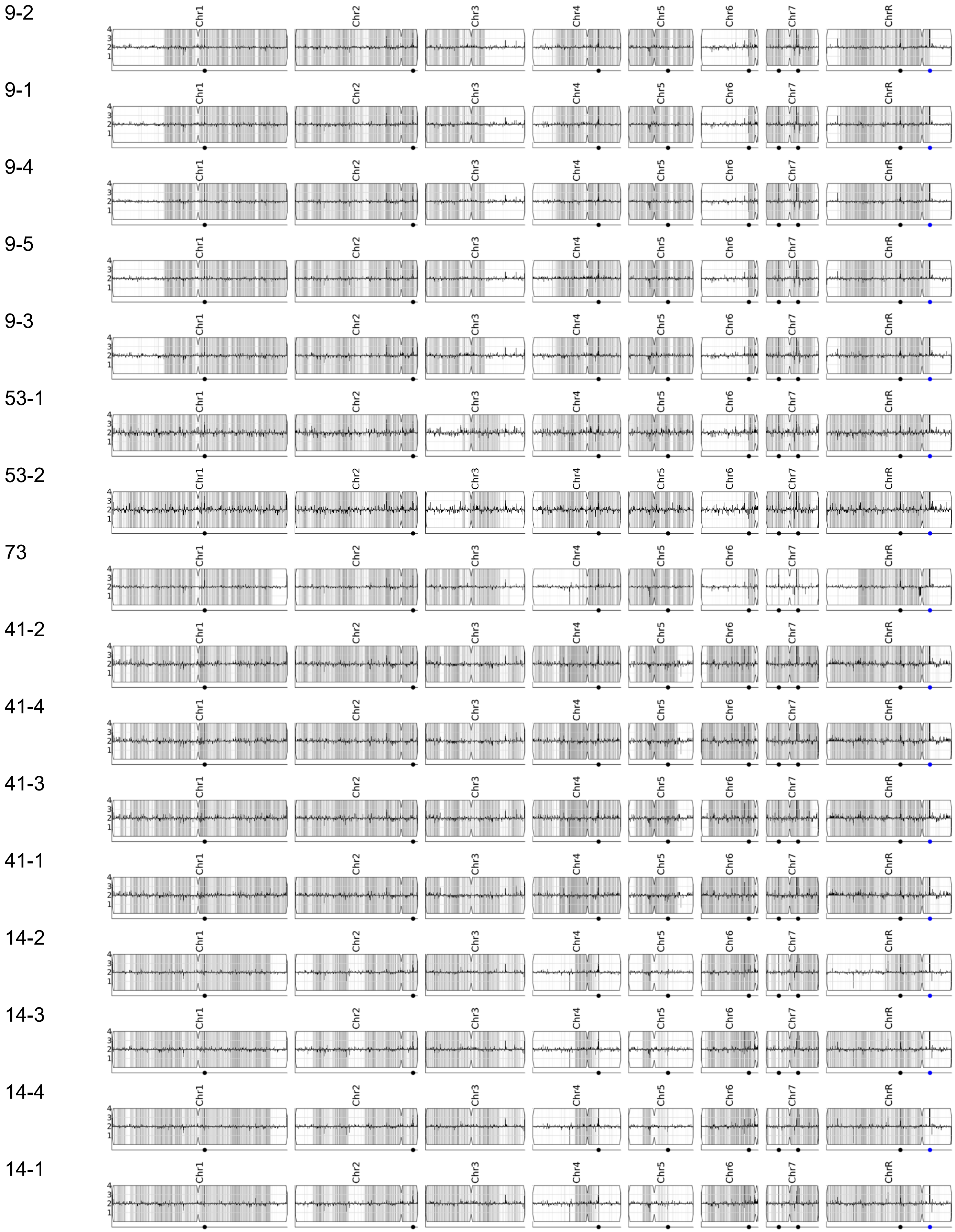

Supplemental Figure S2 (4 of 7)

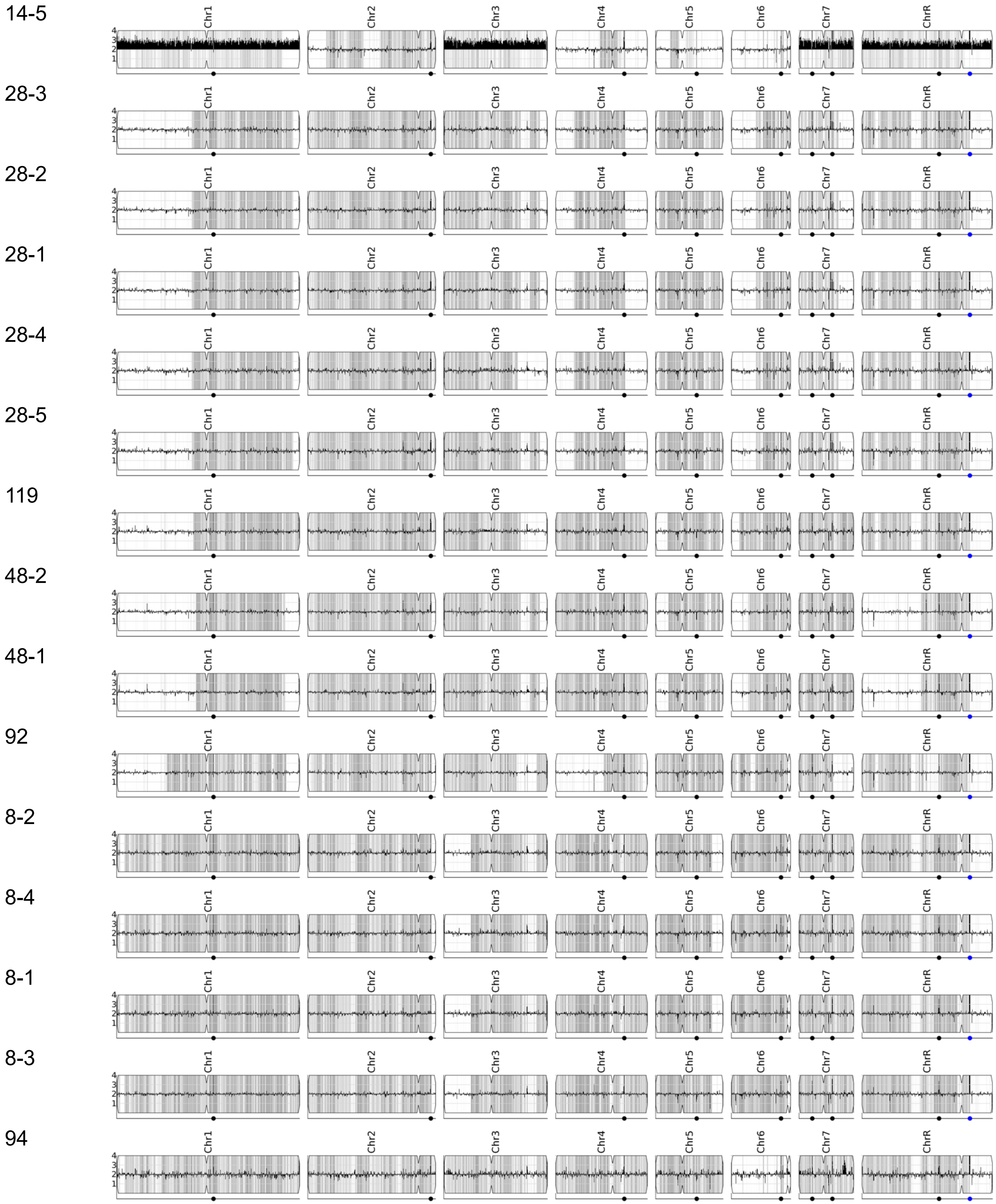

Supplemental Figure S2 (5 of 7)

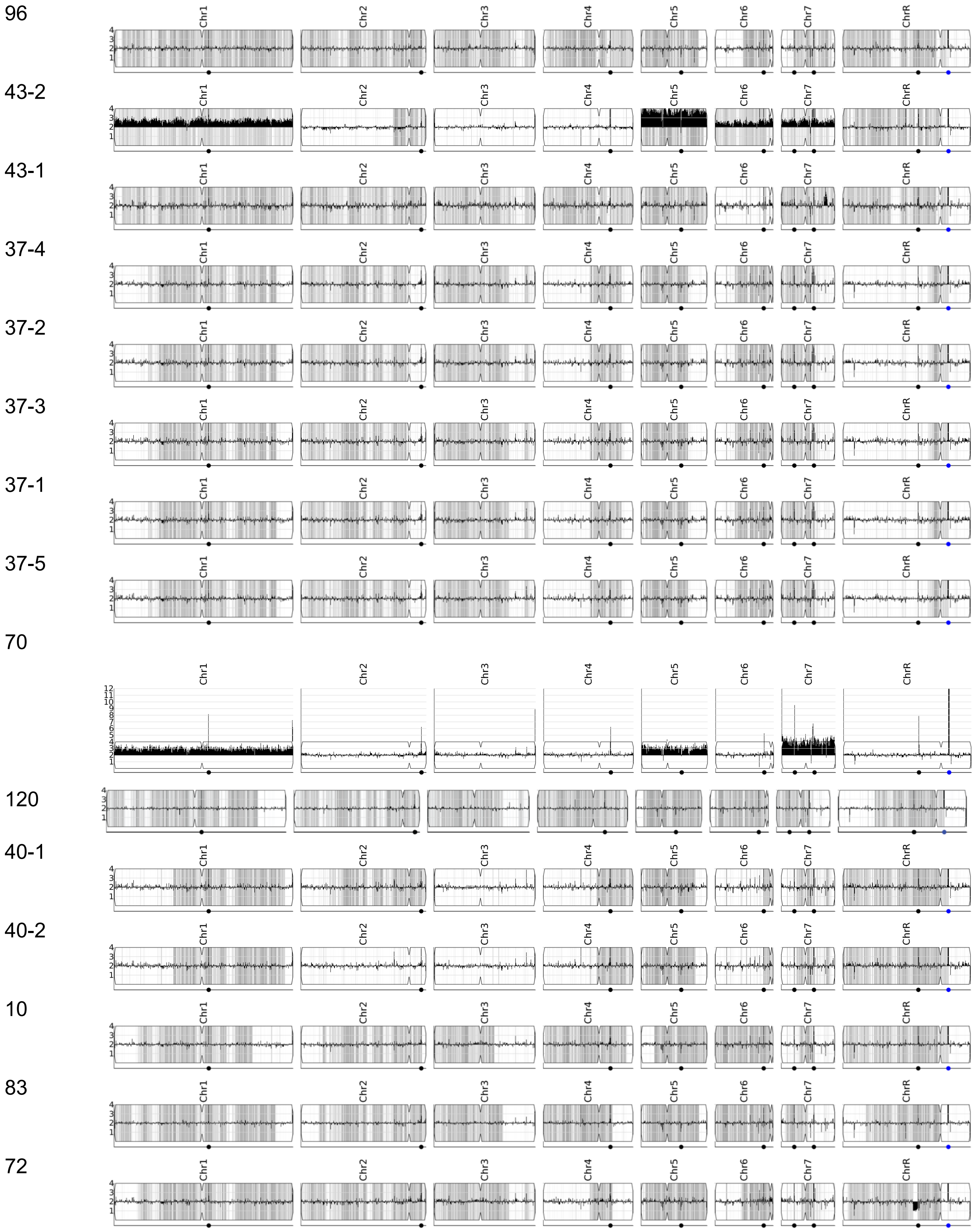

Supplemental Figure S2 (6 of 7)

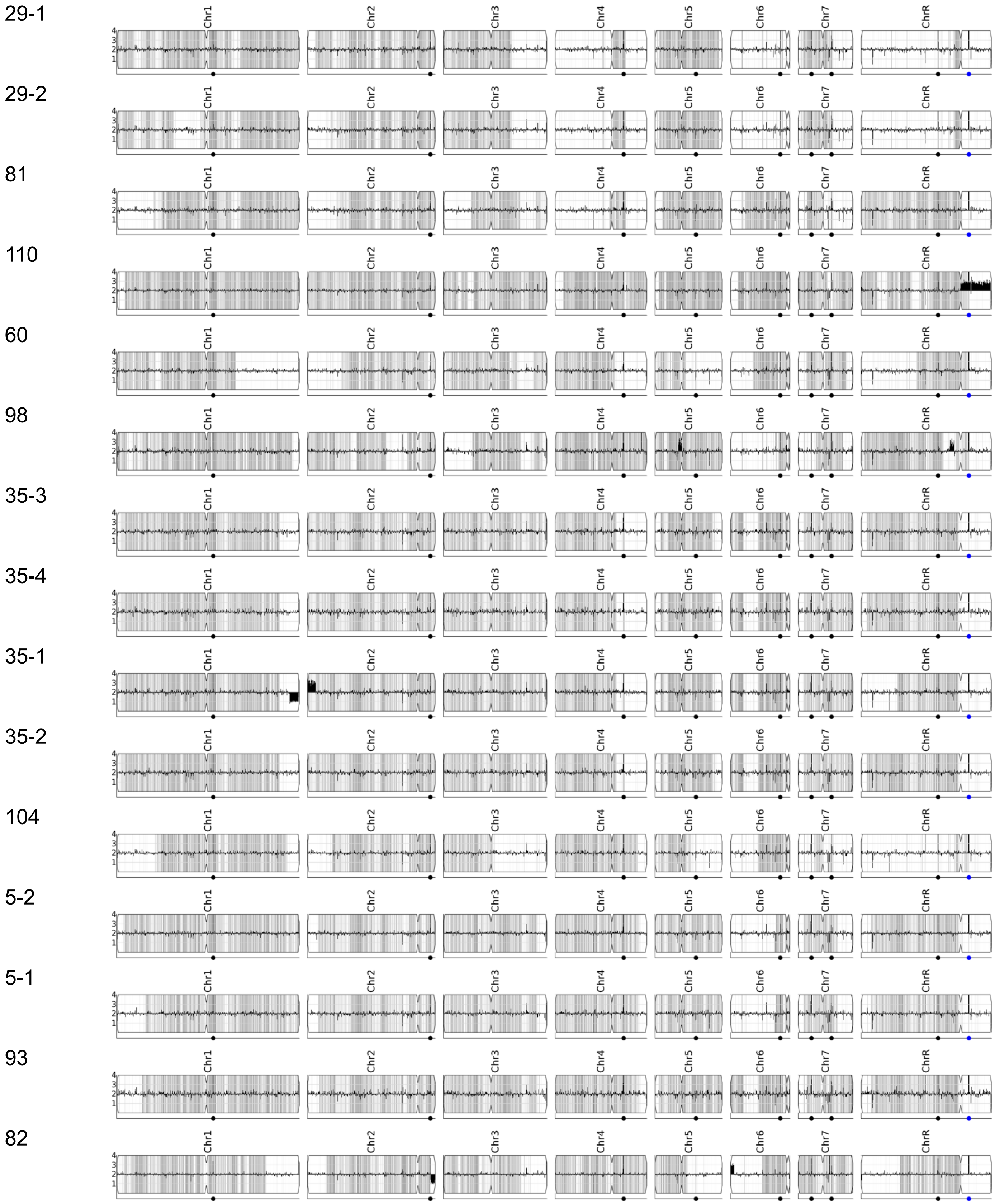

Supplemental Figure S2 (7 of 7)

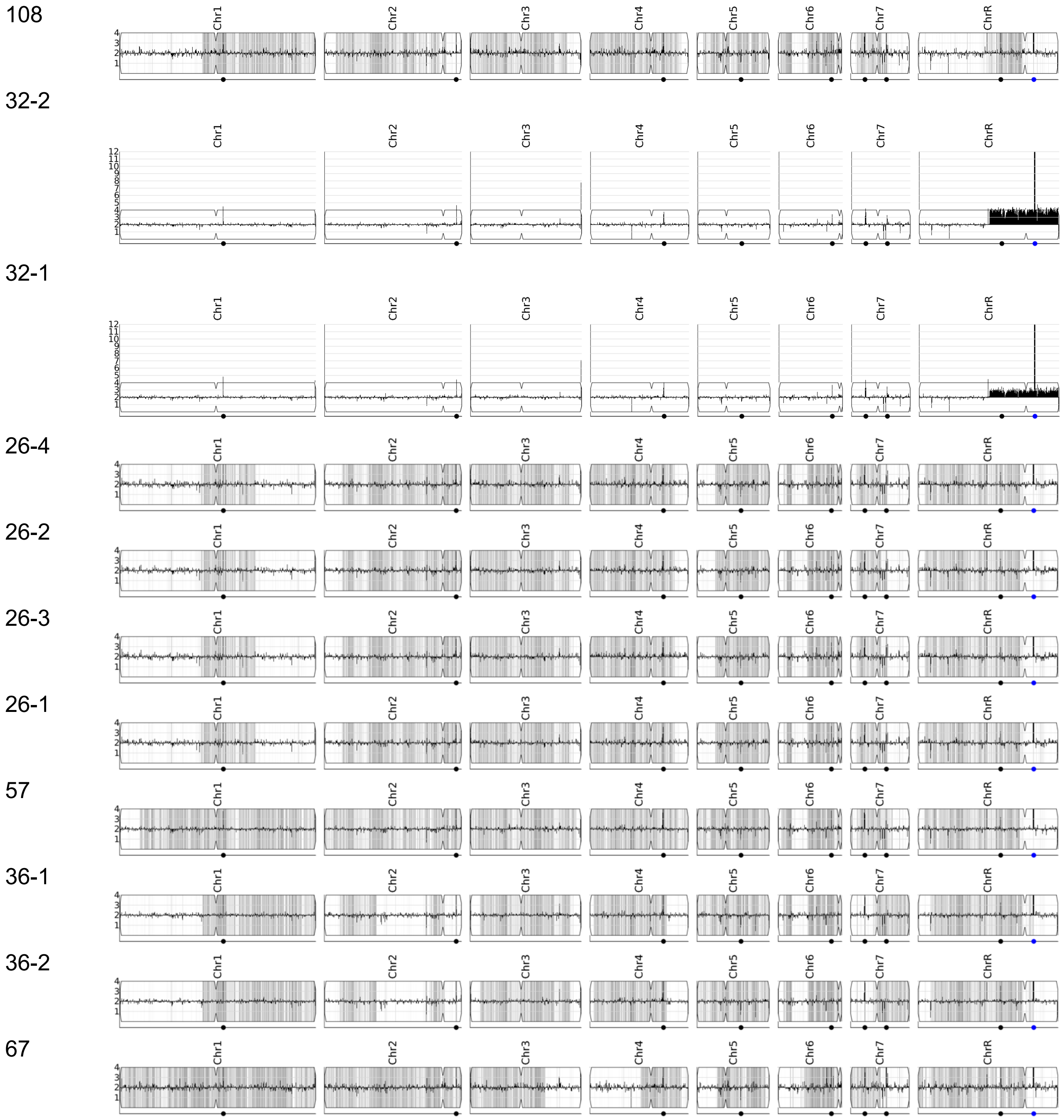


**Supplemental Figure S2 LOH patterns distinguish unrelated strains.** WGS data of 101 study isolates plotted with YMAP. Relative copy number (y-axis) and heterozygous SNP density per 5 kb region in gray (x-axis). Isolates are ordered by the phylogenetic tree in Figure 1.

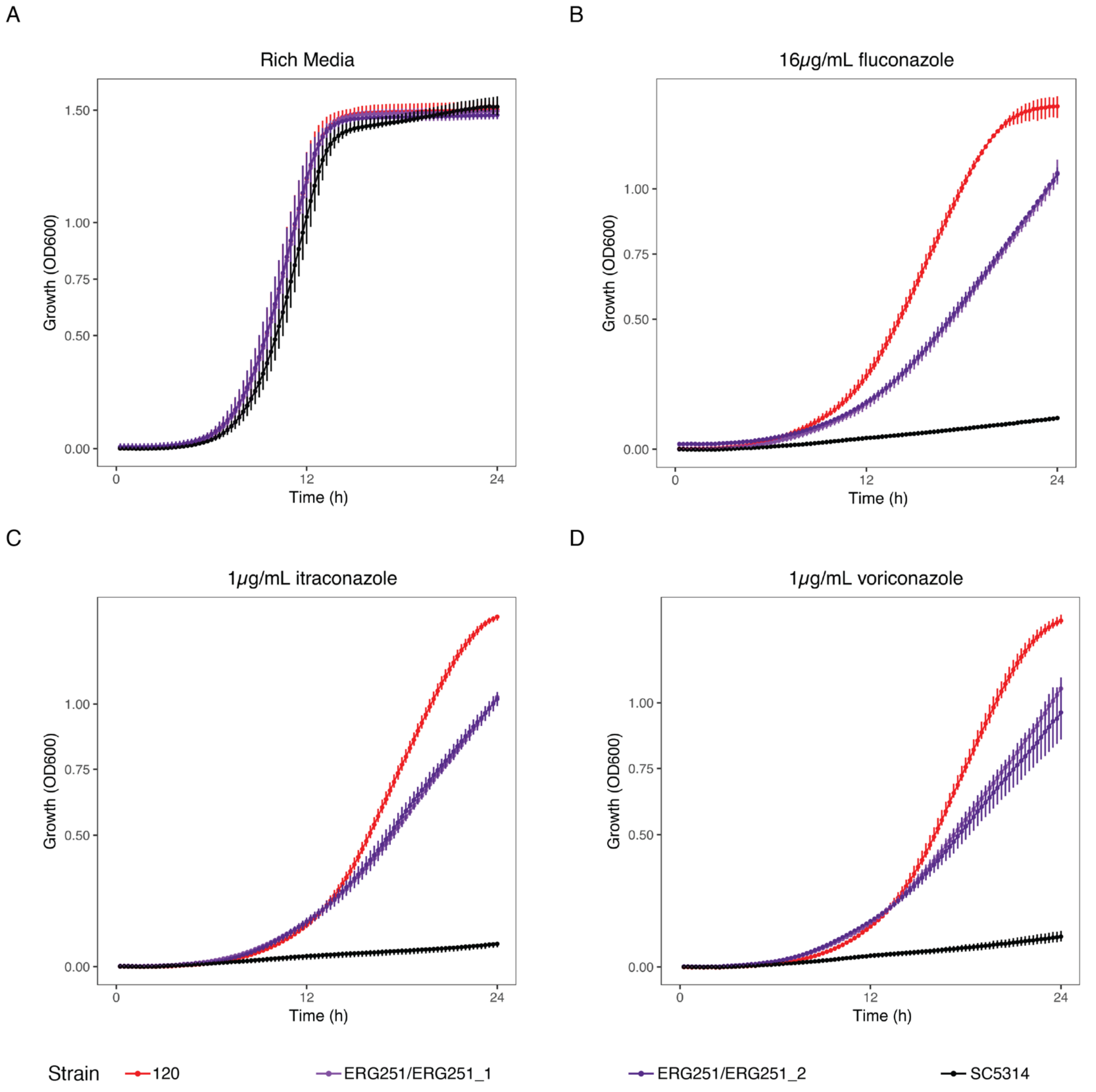


**Supplemental Figure S3 Single allele *ERG251* dysfunction does not have a significant fitness cost and complementation of WT *ERG251* allele increases azole sensitivity.** (A – D) Growth curves of isolate 120 (*ERG251/erg251$^{W145*}$*), two *ERG251/ERG251* engineered strains and reference strain SC5314 (*ERG251/ERG251*). Time in hours on the x-axis and OD600 on the y-axis. (A) Rich media. One-way ANOVA test of AUC-E, p = 0.658. (B) 16ug/mL fluconazole (C) 1ug/mL itraconazole (D) 1ug/mL voriconazole.

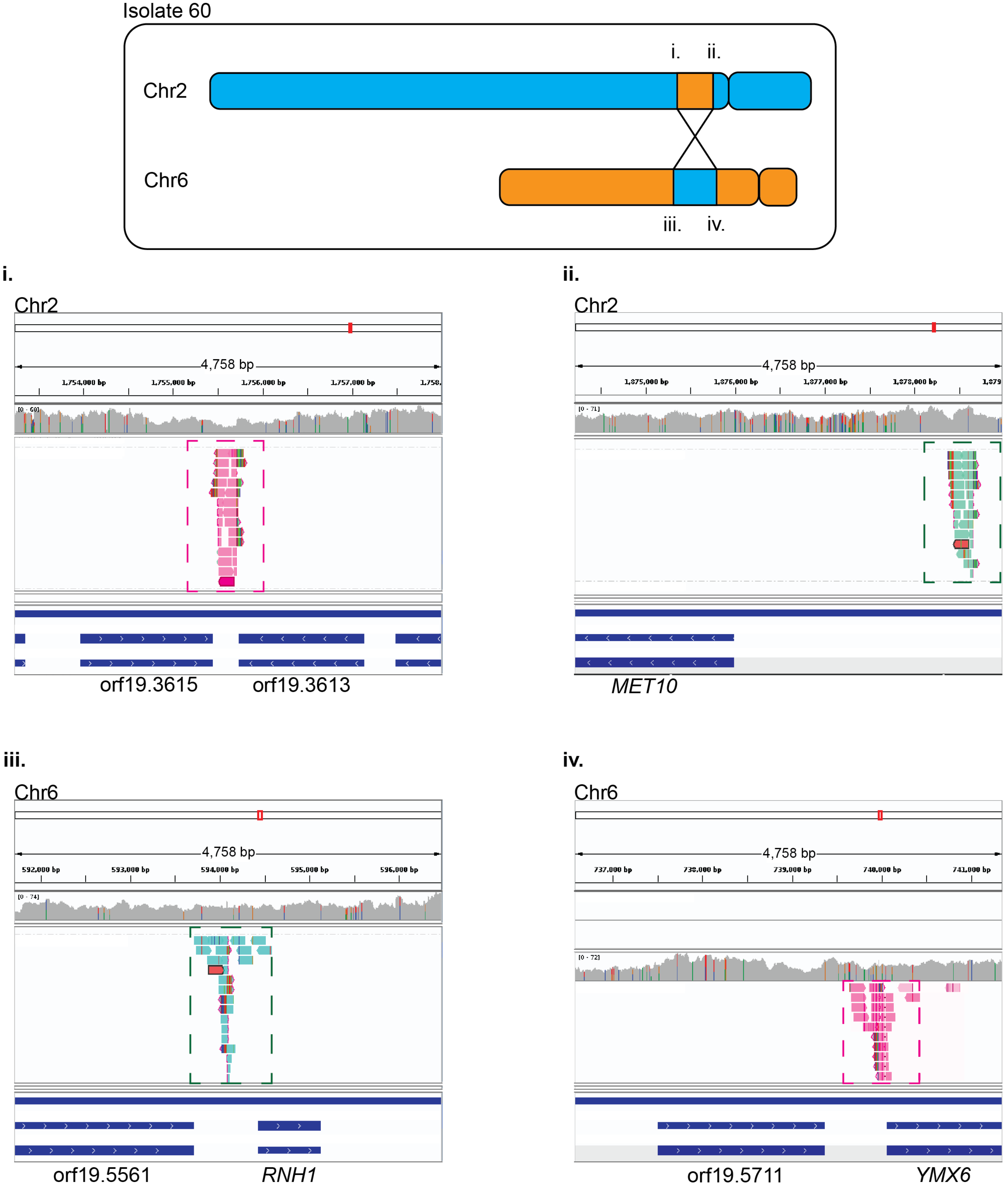


**Supplemental Figure S4 Evidence for reciprocal translocation between Chr2 and Chr6 in isolate 60.** Top: schematic of translocation event. Sequencing read alignments to SC5314 reference genome at the site of the potential translocation, visualized in Integrated Genomics Viewer (IGV). (i and iv) Paired-end analysis of reads mapping at the left side of Chr2 paired to the right side of Chr6 (pink). (ii and iii) Paired-end analysis of reads mapping to the right side of Chr2 and paired to the left side of Chr5 (aqua).

Supplemental Figure S5 (1 of 3)



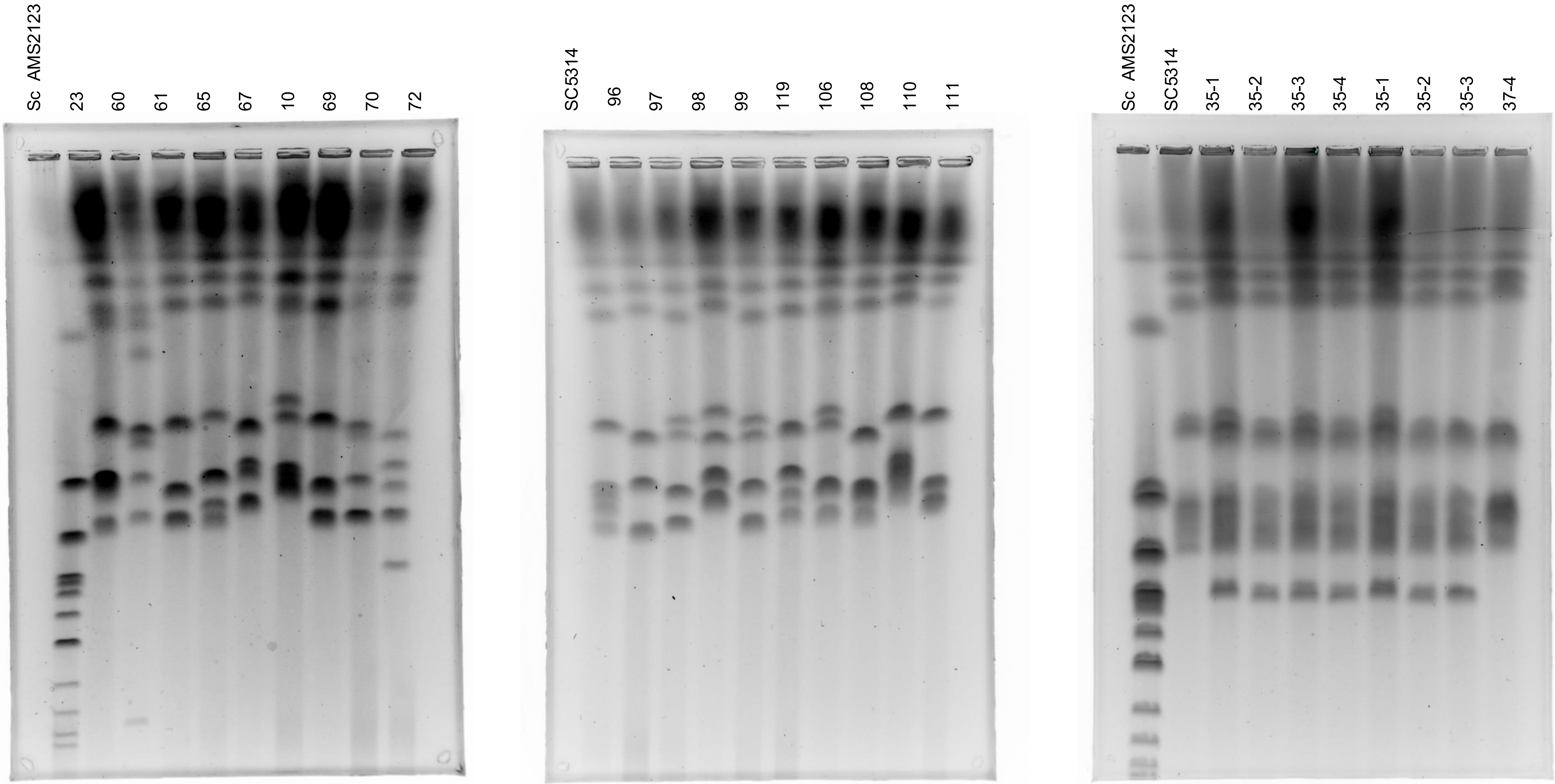

Supplemental Figure S5 (2 of 3)



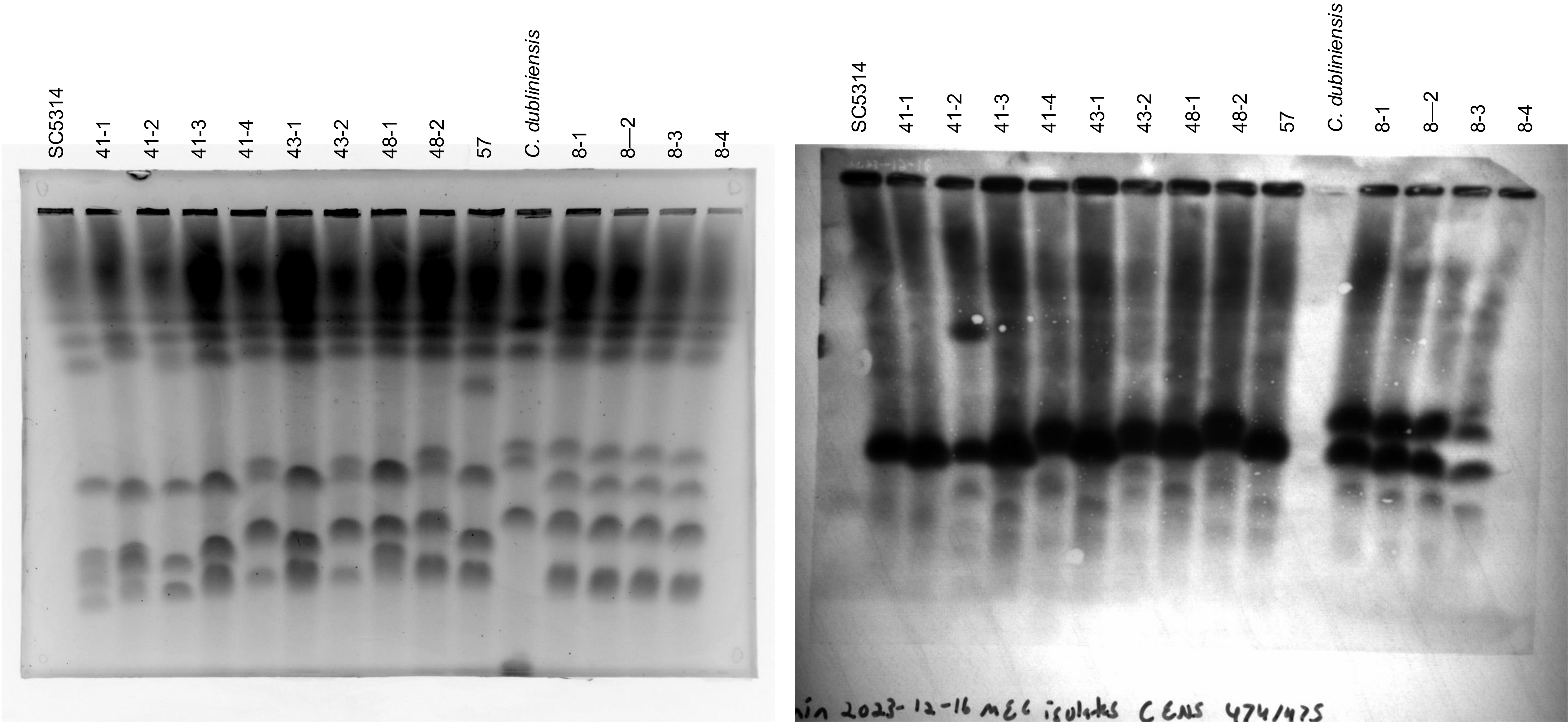

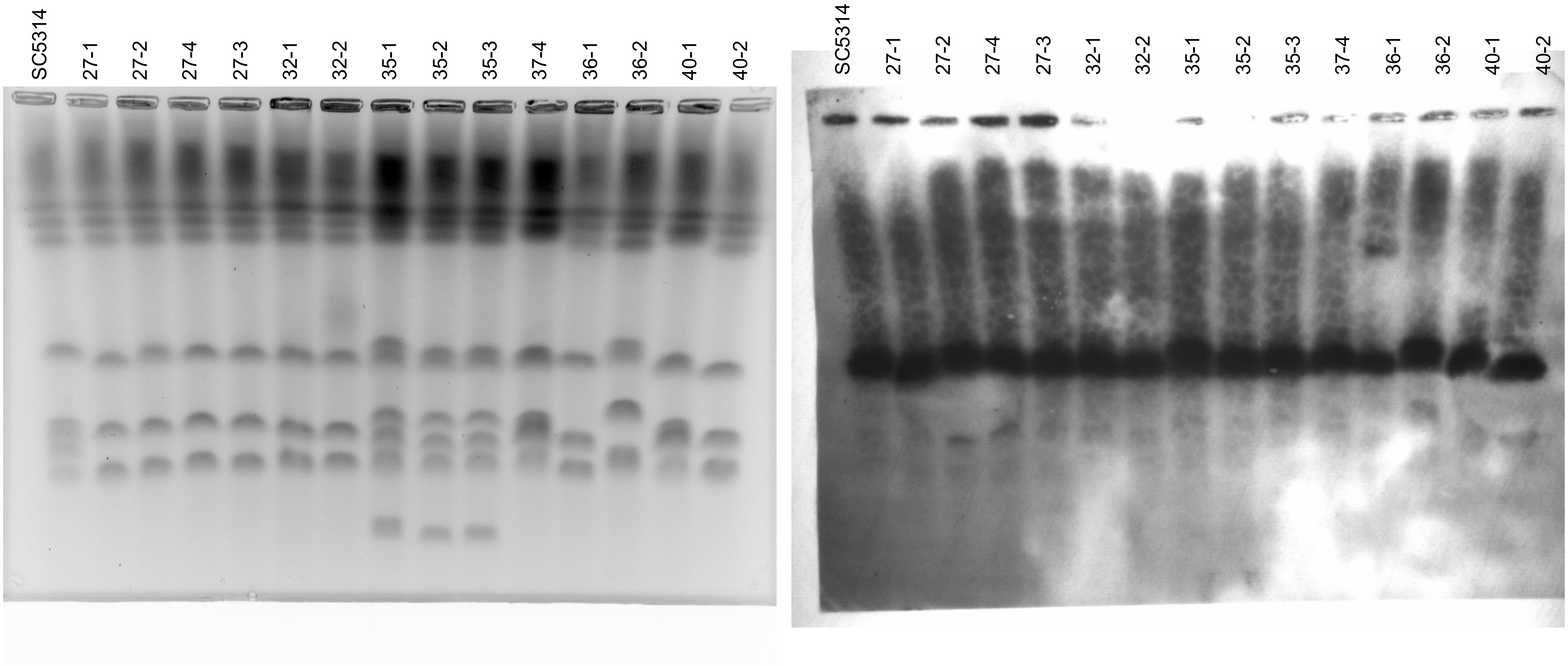


**Supplemental Figure S5 Uncropped karyotype gels and Southern blots**. Full images from Figure 4.

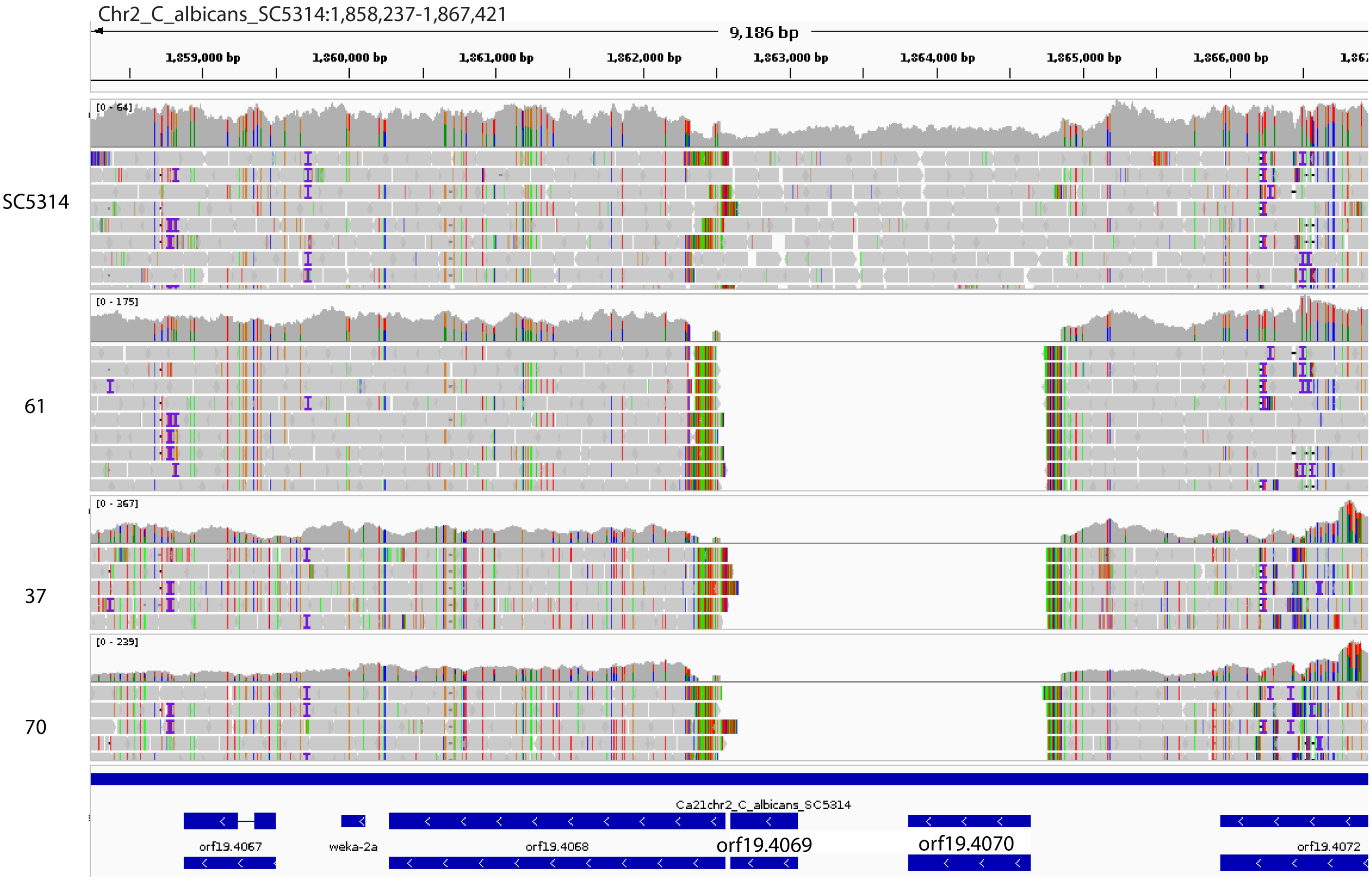


**Supplemental Figure S6 Deletion of orf19.4069 (and dubious orf19.4070) in clinical isolates from multiple phylogenetic clades.** Sequencing read alignment to SC5314 reference genome, visualized in Integrated Genomics Viewer (IGV). For each isolate (labeled at left), sequencing depth is visualized as a coverage histogram above individual read alignments. Colored vertical bars indicate imperfectly mapped reads or indels. No reads map to ORFs19.4069 and 19.4070 in isolates 61, 37, 70. Decreased coverage of SC5314 reads mapped back to the reference genome at ORFs19.4069 and 19.4070 suggest these ORFs could be a heterozygous insertion event.

<u>**Supplemental Data**</u>

Supplemental Table S1: Strain table with MIC, SMG, MLST, clade, genome heterozygosity
Supplemental Table S2: Publicly availabe genomes analyzed in this study
Supplemental Table S3: Pearson's correlation of heterozygosity and phenotypes
Supplemental Table S4: Aneuploid isolate FACS and estimated relative copy number by read depth analysis
Supplemental Table S5: CNV size, breakpoints and associated repeat regions
Supplemental Table S6: Gene amplifications and deletions
Supplemental Table S7: Primers used for WT ERG251 transformation
Supplemental Table S8: CFU counts for Figure 7
